# T cell mediated impairment of epithelial integrity in Crohn’s disease

**DOI:** 10.1101/2024.11.12.621219

**Authors:** Sarah Hamoudi, Nassim Hammoudi, Julie Bonnereau, Anthony Sonn, Elisabeth Capelle, Madeleine Bezault, Victor Chardiny, Joëlle Bonnet, My-Linh Tran Minh, Jean-Marc Gornet, Clotilde Baudry, Hélène Corte, Léon Maggiori, Antoine Toubert, Etienne Becht, Matthieu Allez, Lionel Le Bourhis

## Abstract

T lymphocytes play a major role in intestinal homeostasis, with a particular impact on the balance between self-renewal and differentiation of intestinal epithelial cells (IECs). In Crohn’s disease (CD) patients, the intestinal mucosa is inflamed, epithelium permeability is increased, and IECs present compositional and functional defects. The role of T lymphocytes interactions with IECs in the physiopathology of CD remains in question. Here, we use a three-dimensional human autologous coculture model between purified intestinal organoids derived from primary tissues of CD and non-inflammatory control patients, and mucosal T lymphocytes extracted from the same location. We show that while in homeostatic context T cells support the proliferation and differentiation balance of organoids, mucosal T lymphocytes from CD patients present a high cytotoxicity against IECs. Importantly, this cytotoxicity is a persistent defect overtime in culture. Organoids also show defective intestinal stem cells (ISCs) proliferation and morphological changes. Single cell RNA sequencing after coculture highlights a general response of T cells to the epithelial microenvironment, and more particularly, an increase activation of a pro-inflammatory CD8+ T cells effector population in CD patients compared to controls.

**Graphical abstract:** 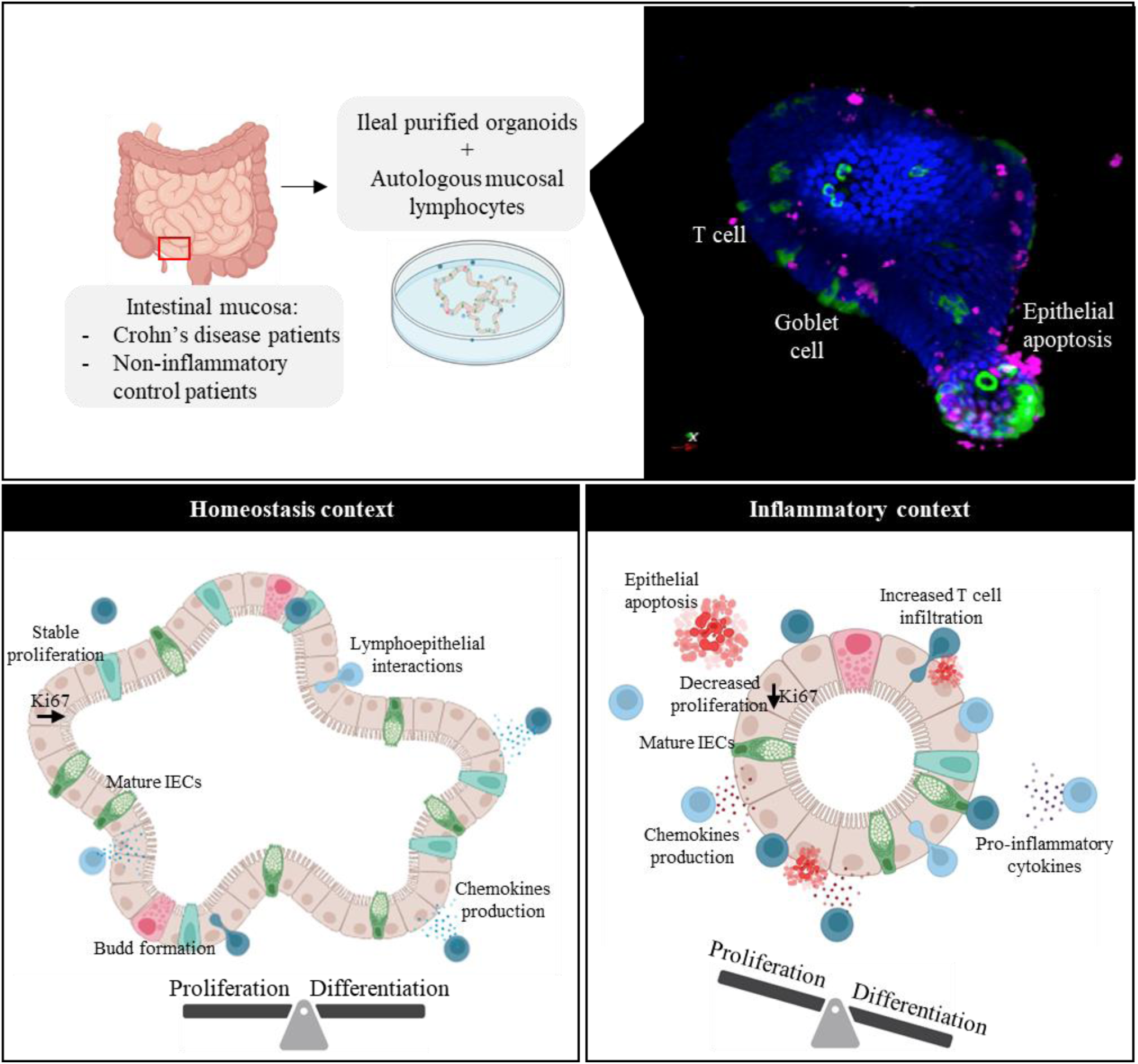

## Introduction

The intestinal epithelium constantly undergoes renewal under the influence of external factors maintaining the proliferation of intestinal stem cells (ISCs) and their differentiation into specialized intestinal epithelial cells (IECs). Defects in the composition and functions of IECs is proposed as a mechanism in the physiopathology of inflammatory bowel diseases, such as Crohn’s disease (CD). CD is characterized by an inflamed intestinal mucosa, triggered by microbial translocation leading to exacerbated immune responses in genetically predisposed patients.^1,2^ Differentiation and function of epithelial cells, such as Paneth cells, which produce anti-microbial peptides, are impaired in these diseases.^3,4^

Among factors impacting IEC biology, T cells of the intestinal mucosa are known to influence the self-renewal and differentiation balance of IECs. These so-called resident memory T cells (Trm) have a key localization at the interface between the host and its environment and are found in very large numbers in the human ileal mucosa^5,6^ and thus, may have a pivotal role in CD. Altered lymphoepithelial interactions may be involved in the pathophysiology of CD as defects in interactions between epithelial cells and peripheral T cells are demonstrated in IBD patients through coculture model showing overstimulation of CD4+ T cells.^7^

A number of studies have thus focused on determining the cytotoxic feature of mucosal T lymphocytes in CD. Clonal expansions of persistent pro-inflammatory NKG2D+ CD4+ T lymphocytes subset in the inflamed mucosa of CD patients participating to epithelium damage have been highlighted.^8^ This observation is supported by other patient studies that highlight an increased subset of resident memory T cells in the *lamina propria* of CD patients.^9^ This same population is reported to be associated with disease relapse and enriched in Th17 and Th1 cells producing pro-inflammatory cytokines.^9,10^ The role of CD8+ Trm cells is currently still debated. Study of TCR repertoire in the ileal mucosa of CD patients shows clonal expansions in CD8+ T cells reported to be associated with post-operative recurrence.^11^ We highlighted two functionally different subpopulations, expressing CD103 or KLRG1 molecules at their surface, showing altered proportions in CD.^12^ Hence, Trm cells are implicated in CD pathogenesis, but the mechanisms involved and their potential as treatment targets remains to be fully elucidated.

In recent years, the rise of organoid models has facilitated the study of stem cells and organ development, but also the study of specific questions about relations between different cell types and their involvement in pathologies.^13^,^14^ Intestinal organoids have thus emerged as great tools for IBD research. Organoids from IBD patients keep the characteristics from the tissue of origin and are used to compare morphological and phenotypic changes, highlighting defects related to the pathology. Assessment of organoid formation efficiency, budding, growth, maturation, but also gene analysis, describes significant changes associated with the disease; and are used to test the efficiency of various therapeutic approaches.^15,16,17^

A number of studies using intestinal organoids have focused on the crosstalk with T cells and demonstrate their importance in homeostasis. Indeed, the balance between stem cell proliferation and differentiation involves indirect interactions with the production of cytokines by T cells, as regulatory mediators support ISCs proliferation and other rather pro-inflammatory cytokines induce differentiation.^18,19^ Return to homeostasis after epithelial damage is also reported to depend on cytokine production by T cells, as demonstrated by the link between IECs stimulation with IL17A leading to secretory lineages differentiation.^20^ However, other studies demonstrate the harmful potential of T cells, such as the cytotoxic effect of IFNγ produced by T cells in inflammatory context leading to ISCs depletion.^21^ A more recent study from our team also highlights the cytotoxic potential of autologous tissue-resident memory T lymphocytes from CD patient against organoids involving direct interactions through NKG2D and CD103 but also through cytokine production.^22^ The use of human organoid models with immune cells in autologous conditions therefore enables not only the study of fundamental questions, but also the test of therapeutic compounds in a wide range of pathologies such as auto-immune and inflammatory diseases or cancers.^23,24^

In this study, our aim was to evaluate the impact of mucosal lymphocytes on IEC differentiation and survival in CD patients. We used an innovative model of purified small intestinal organoids from patients cocultured with autologous resident memory T cells (Trm). Purified organoids were obtained by repeated passages leading to undifferentiated organoids composed almost exclusively of ISCs. We showed that in homeostatic context Trm supported ISCs proliferation and epithelial maturation while preserving organoid morphology. However, with cells from CD patients, Trm impacted the proliferative capacity and morphology of organoids while inducing epithelial cell death. Transcriptomic analysis at the single cell level of T cells post-coculture showed changes in T cell gene signatures in response to the epithelial microenvironment. Interestingly the emergence of an effector memory CD8+ T cells population with a pro-inflammatory activated Th1 profile in T cells from CD patients was observed in coculture with autologous organoids but not in non-inflammatory control cocultures.

## Results

### An autologous coculture system to study relationship between purified intestinal organoids and long-term cultured mucosal T lymphocytes

We first characterized ileal and colon derived organoids after several passages to obtain structures mostly composed of ISCs. In bright field microscopy, both ileal and colon organoids adopted a hallow bubble-like shape, with a thin membrane corresponding to a monocellular epithelial layer. They were polarized, with the apical face turned towards the inside, representing the intestinal lumen, and the basal face turned towards the outside, directly accessible. After a few days of culture, buds appeared, and even more so, in ileal organoids that can adopt an enriched crypt-villi like structure. Over time, their interior darkens, filling with senescent differentiated cells shedding into the lumen. (Fig1.B)

**Fig 1:**
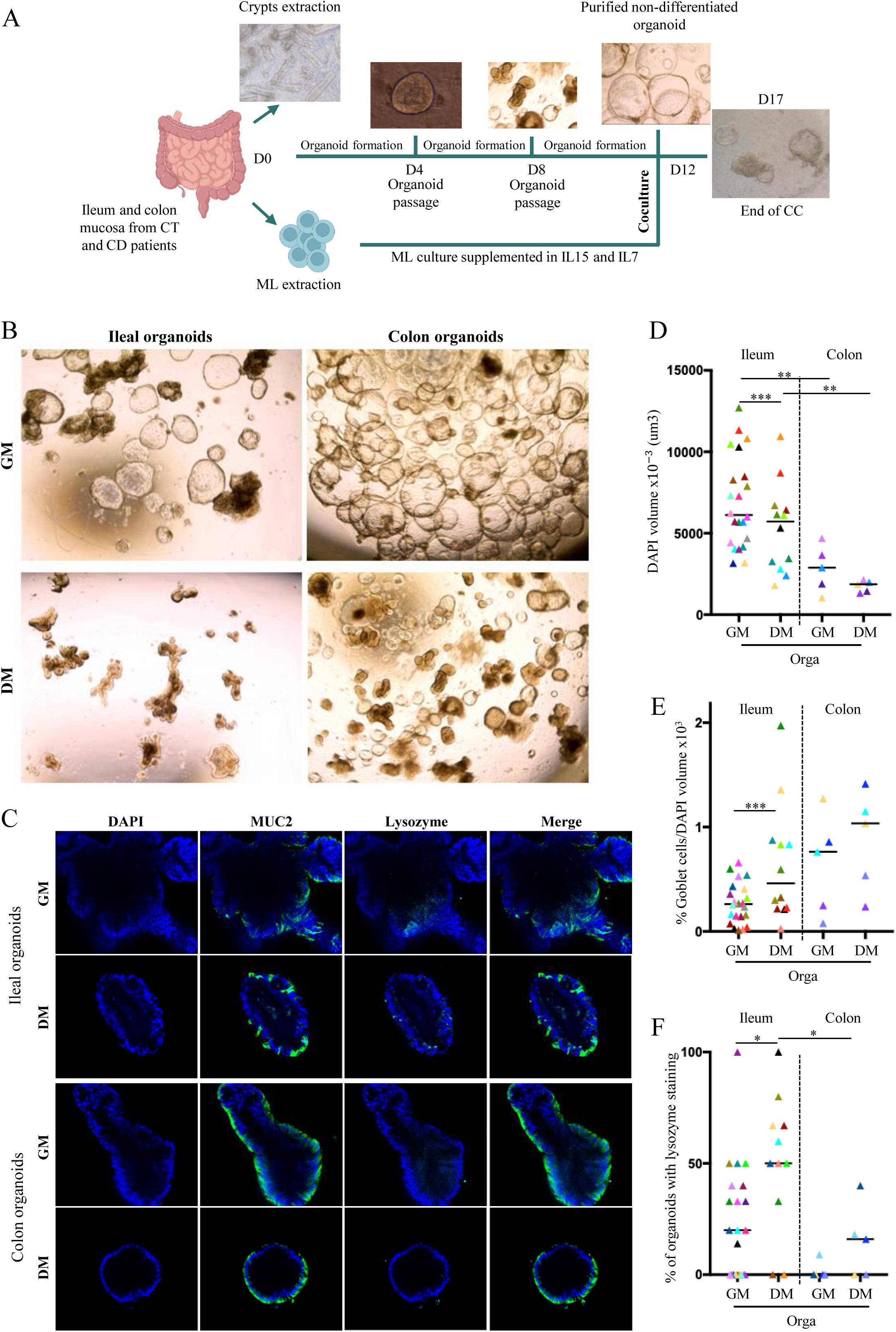
Characterization of human ileal and colon organoids. (A) Experimental scheme of autologous cocultures (B) Observation with bright field microscopy (20X) of organoids in GM or DM after 5 days of growth (C) Observation with confocal microscopy of organoids in GM or DM (200x) (D) Quantification of organoids size through DAPI volume staining after 5 days in GM or DM (E) Quantification of the percentage of Goblet cells over the DAPI volume after 5 days in GM or DM (F) Quantification of the percentage of organoids with lysozyme after 5 days in GM or DM Each point corresponds to the median of the values obtained for organoids of a given patient. GM: Growth medium, DM: Differentiation medium, Orga: Organoids alone. Ileum: in DM n=12, in GM n=22; Colon: n=5

After five days in medium supporting growth (GM) or inducing differentiation (DM), we compared ileal and colon organoids size and identified the cell types that constitute them by confocal microscopy. (Fig1.C) Colon organoids were about twice as small as those from ileum. Inducing differentiation with the specific medium also decreased organoid size. (Fig1.D) Assessment of their differentiation capacity showed that in GM colon organoids differentiated more towards Goblet cells than those from ileum (around three times more), using Mucin2 (Muc2) specific staining. Inducing differentiation led to increase Goblet cell numbers in ileal organoids. This effect was not observed in colon organoids as they already had a high number of Goblet cells. (Fig1.E) Lysozyme staining was used to evaluate the presence of Paneth cells. Few ileal organoids in GM produced lysozyme but DM increased significantly this number. Colon organoids produced almost no lysozyme even in DM. (Fig1.F) Taken together, these observations corroborate current knowledge on intestinal physiology as more Goblet cells are found in the colon compared to ileum, where classically Paneth cells are found.

### Autologous mucosal T lymphocytes support epithelial maturation by inducing differentiation at homeostasis

We first checked the viability and phenotype of resting mucosal lymphocytes (ML) by flow cytometry before coculture. Survival was assessed by DAPI staining and no difference was observed between ML isolated from ileum or colon, from control and CD patients. Ileum derived T lymphocytes showed three times more CD4+ T cells than CD8+ T cells in CD patients whereas a comparable proportion was found in control patients. Colon ML showed similar phenotype whether from control or CD patients. The proportion of CD8+ T cells on CD4+ T cells was similar, and the expression of ligands and receptors of interest were comparable. (Supp1)

We then tested the impact of autologous ML on the maturation of ileal organoids derived from control patients. Confocal microscopy quantification showed that ML from control patients did not decreased organoid size in GM but did in DM. (Fig2.A,B) Increase differentiation was observed in cocultures compared to organoids alone as percentages of Goblet cells and lysozyme staining augmented significantly. (Fig2.C,D) As organoid size stayed similar in GM it can be concluded that in homeostatic context autologous mucosal lymphocytes increased IECs differentiation towards secretory lineages.

**Fig 2:**
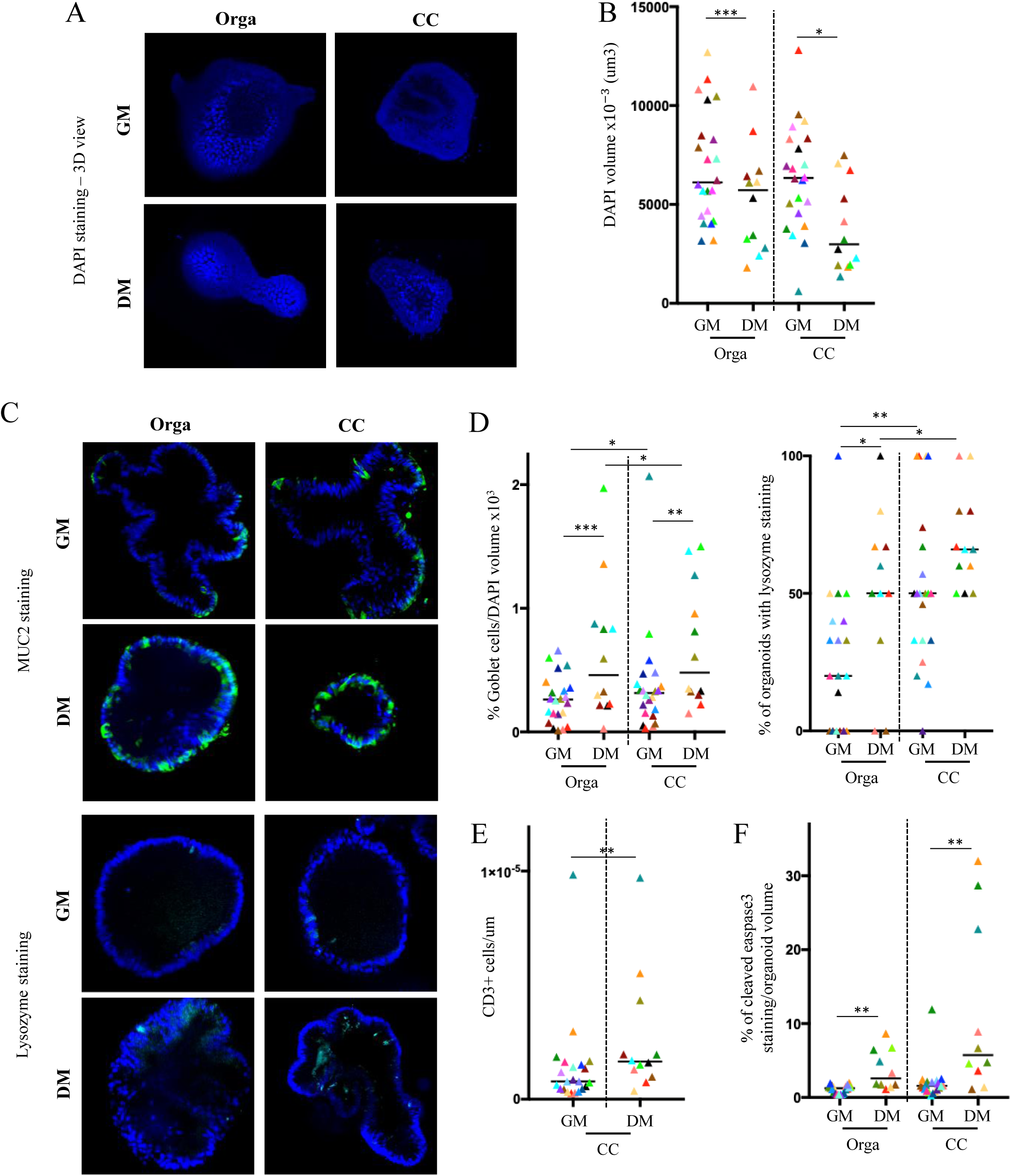
Mucosal T lymphocytes support epithelial maturation at homeostasis. (A) 3D view of ileal organoids alone or in coculture with T cells stain with DAPI (blue) after growing 5 days in GM or DM by confocal microscopy (200X) (B) Quantification of ileal organoids size alone or cocultured with T cells through DAPI volume staining (C) Confocal microscopy (200X) of ileal organoids alone or cocultured with T cells in GM or DM (Muc2: green; lysozyme: cyan) (D) Quantification of percentage of Goblet cells over DAPI volume and organoids with lysozyme in ileal organoids alone or in coculture (E) T cell infiltration assessed with the number of CD3+ T cells in contact over DAPI volume (F) Organoid epithelial cell death assessed with the cleaved Caspase-3 fluorescence volume over DAPI volume GM: Growth medium; DM: Differentiation medium; CC: cocultures, Orga: organoids alone. Each point corresponds to the median of the values obtained for organoids of a given patient. In DM n=12, in GM n=22

The lymphocyte infiltration rate and epithelial cell death in cocultures were also evaluated. Epithelial cell death was evaluated through the cleaved caspase-3 staining normalized on organoid size. Number of CD3+ cells in close contact with organoids were counted over organoid size to obtain a number of T cells interacting with organoids per µm. When differentiation was induced by the medium (DM), the ML infiltrated ileal organoids more. (Fig2.E) These organoids were more mature, but they were also smaller. Cleaved caspase-3 quantification showed very low IEC death in GM but a significant increase in DM. Autologous ML did not increase epithelial cell death in control ileal organoids. (Fig2.F) These observations showed that ML from control patients support epithelial maturation without inducing aberrant epithelial cell death.

### Autologous ML from CD patients change the morphology of purified ileal organoids and impact their proliferative capacity

We then wanted to assess if autologous ML from CD patients had a different impact on organoids maturation. First, ileal CD organoids alone were compared to those from control patients. (Fig3.A) DAPI quantification showed that ileal organoids from CD patients were the same size than those from controls. (Fig3.B) They also tended to have a slightly higher percentage of Goblet cells, and more organoids produced lysozyme. (Fig3.C,D) Quantification of budding organoids percentage for each patient revealed a similar number of budding organoids in control and CD patients. (Fig3.E) Based on these parameters, morphology and maturation of ileal organoids alone from control and CD patients were comparable.

**Fig 3:**
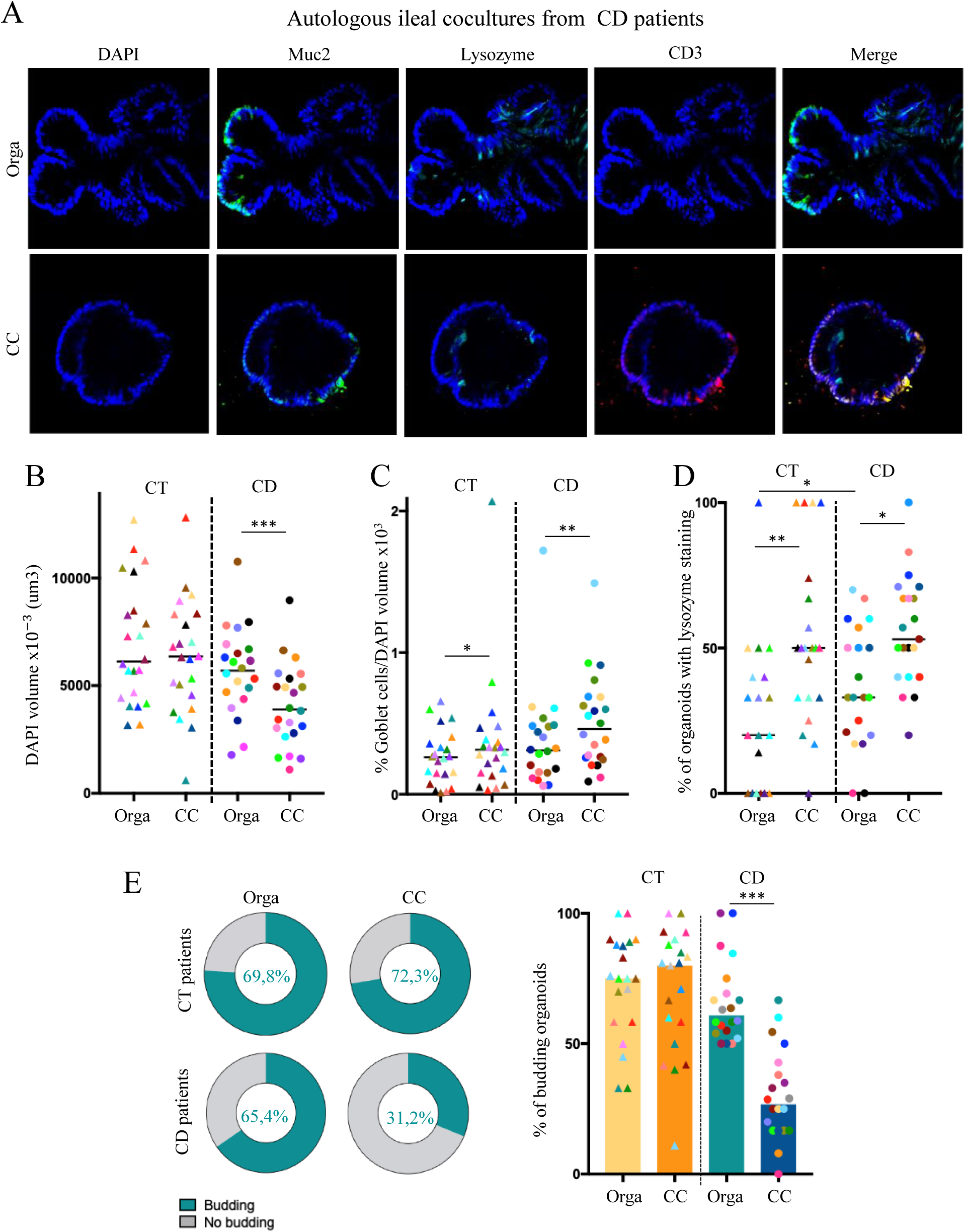
Morphological changes suggest proliferation impairment in cocultured ileal organoids from CD patients. (A) Confocal microscopy (200X) of autologous ileal coculture from CD patients (DAPI: blue, Muc2: green, lysozyme: cyan, CD3+ cells: red) (B) Quantification of ileal organoids size through DAPI volume staining (C) Quantification of the percentage of Goblet cells over the DAPI volume (D) Quantification of percentage of organoids with lysozyme (E) Quantification of the percentage of budding organoids from control or CD patient, alone or in coculture CC: cocultures, Orga: organoids alone CT: control patients, CD: Crohn’s disease patients. Each point corresponds to the median of the values obtained for organoids of a given patient. CT patients n=21, CD patients n=22

Based on the same parameters, CD patients’ cocultures were compared with those from control patients. (Fig3.A) While ML from controls patients had no effect on organoids size, adding autologous ML with CD organoids significantly decreased their size. (Fig3.B) Quantification of Goblet cells and organoids producing lysozyme showed in CD organoids cocultures an increased ratio of differentiation towards secretory lineages. (Fig3.C,D) Moreover, the percentage of budding organoids per patient revealed that organoids from CD patients in coculture were less budding than alone but also less than those from control patients in cocultures. (Fig3.E) These observations showed immune mediated morphological changes in CD patients organoids and suggested that autologous ML from CD patients may impact their proliferative capacity.

### Autologous ML from CD patients disrupt the proliferation/differentiation balance of ileal organoids

To test whether the MLs had an impact on the organoid proliferative potential, quantification of the percentage of Ki67+ live proliferative cells by flow cytometry was performed. In control cocultures, the percentage of Ki67+ live IECs remained comparable to that from control organoids alone. Although the percentage of Ki67+ proliferative cells in CD organoids alone was also comparable to that of control organoids, the addition of autologous MLs led to a significant decrease of the Ki67+ proliferative population. (Fig4.C) This result was then confirmed by quantifying the proliferative level of organoids by confocal microscopy (quantification of Ki67+ volume staining reported to DAPI volume staining). The same pattern was observed: only the proliferation rate of cocultured CD organoids was decreased. (Fig4.D) The quantification of expression level of stem-related genes Ki67 and Sox9 in the same conditions showed a slight decrease in control cocultures, but a significant decrease of Sox9 expression was observed in CD cocultures. (Fig4.E) Taken together, these results demonstrated that autologous mucosal lymphocytes from CD patients negatively impacted the IECs proliferative capacity while those from controls maintained the proliferation/differentiation balance.

**Fig 4:**
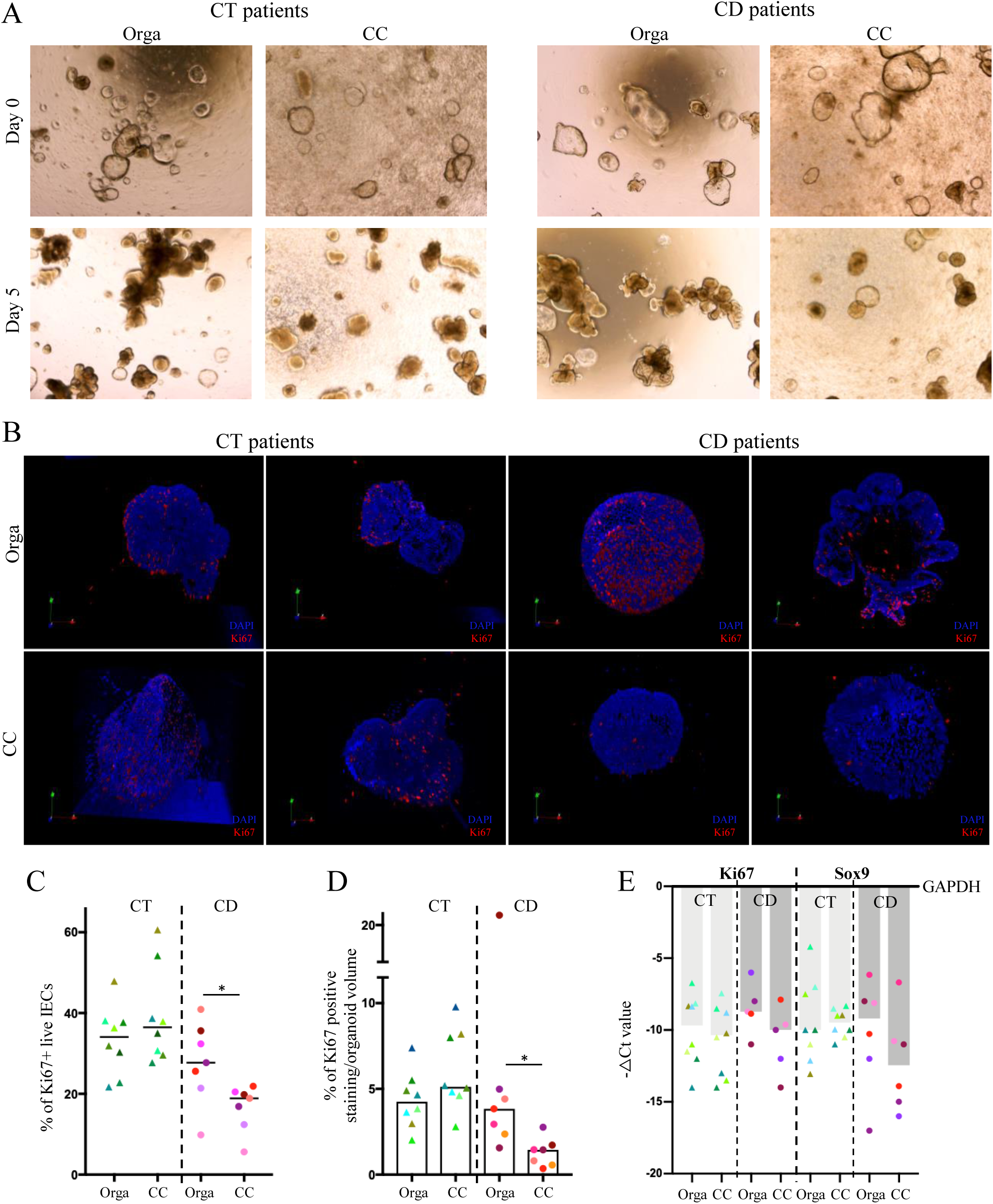
Autologous ML disrupt the proliferation/differentiation balance of CD patient ileal organoids. (A) Observation with bright field microscopy (20X) of ileal organoids morphology evolution after 5 days of growth (B) Observation with confocal microscopy (200X) of ileal organoids morphology evolution after 5 days of growth (C) Quantification by flow cytometry of the percentage of proliferative Ki67+ live IECs (D) Quantification of the organoid proliferation level through Ki67 volume staining over DAPI volume (E) Gene expression level of Ki67 and Sox9 in ileal organoids Orga: organoids, CC: coculture, CT: control patients, CD: Crohn’s disease patients. Each point corresponds to the median of the values obtained for organoids of a given patient. CT patients n=8, CD patients n=7

### Immune mediated epithelial cytotoxicity is a persistent defect in CD patients cocultures

We next assessed the T cell-mediated cytotoxicity from CD patients in ileal organoids after several passages and with long-term lymphocyte culture. (Fig5.A) Lymphocyte infiltration quantification showed that ML from CD patients interacted more with ileal organoids than those from control patients, although attracting signals were produced in both cases. (Fig5.B) Moreover, these contacts were accompanied by an increased epithelial cell death only in CD cocultures. (Fig5.C) These results show that purified organoids passaged several times derived from the ileum of CD patients are susceptible to an autologous T cell mediated cytotoxicity which is not observed in control autologous cocultures.

**Fig 5:**
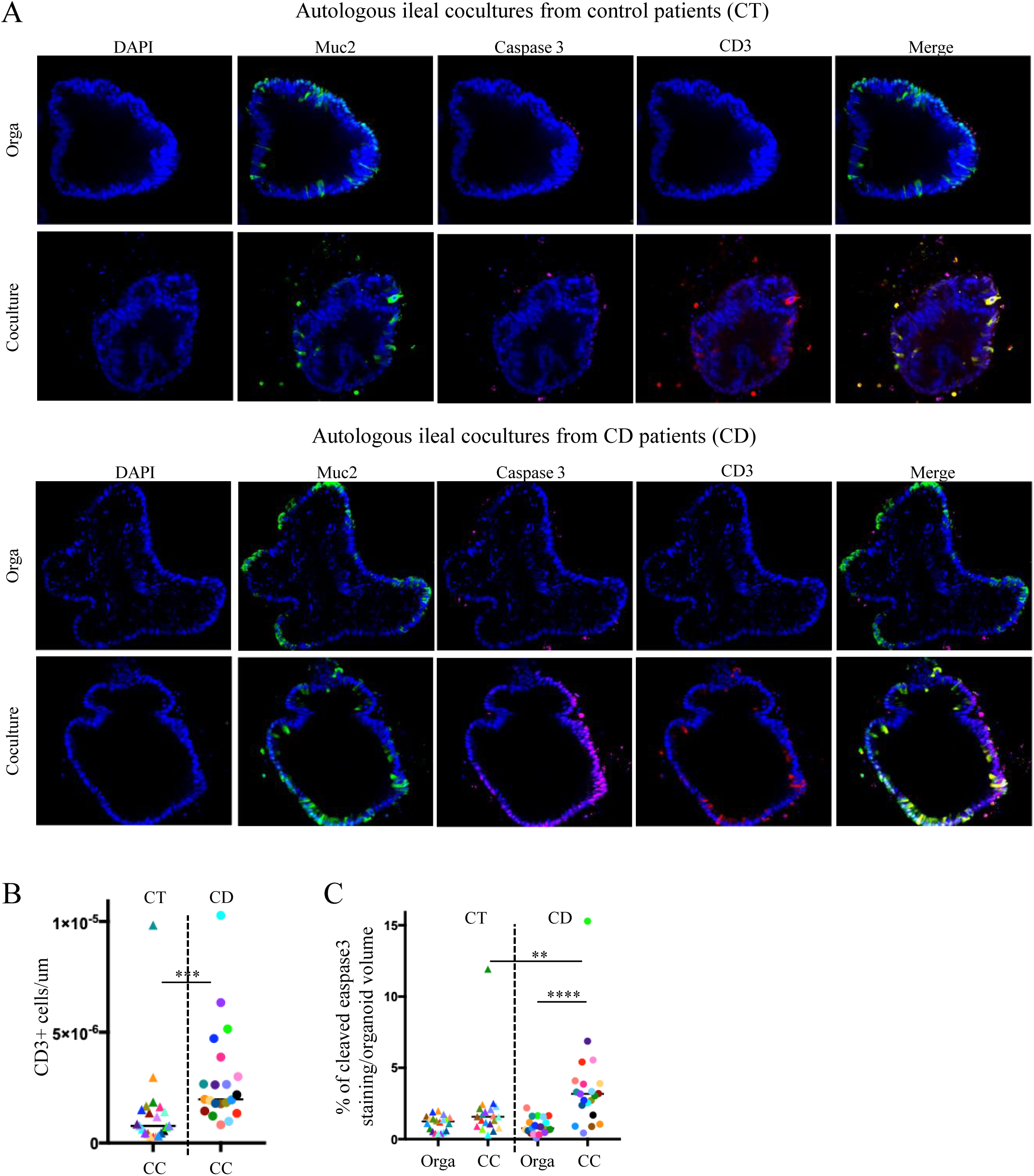
Immune mediated epithelial cytotoxicity in CD patients cocultures is a persistent defect. (A) Confocal microscopy (200X) of autologous ileal cocultures from control and CD patients (DAPI: blue, Muc2: green, Caspase3: pink, CD3+ cells: red) (B) T cell infiltration assessed with the number of CD3+ T cells in contact over DAPI volume (C) Organoid epithelial cell death assessed with the cleaved Caspase-3 fluorescence volume over DAPI volume All experiments were made in GM; CT: control patients; CD: CD patients; CC: cocultures, Orga: organoids alone. Each point corresponds to the median of the values obtained for organoids of a given patient. CT patients n=20, CD patients n=21

### The sensitivity of IECs to T lymphocytes from Crohn’s disease patients is not an epithelial defect

We then questioned whether the proliferative defect and epithelial cytotoxicity observed were consequences of T cell functions or if the IECs from CD patients presented an increased intrinsic sensitivity. We hence designed an allogenic model of coculture between ileal organoids from CD or control patients and T cells sorted from PBMCs of healthy donors. (Fig6.A) As expected, quantification of epithelial cell death by confocal microscopy showed increase caspase-3 cleavage in organoids from control patients cultured with donor T cells resulting from allogenic response. However, no epithelial cell death was induced in CD patient organoids cocultured with peripheral T cells from healthy donors, suggesting, on the contrary to what we hypothesized, a desensitization of IECs to this allogenic response. (Fig6.B)

**Fig 6:**
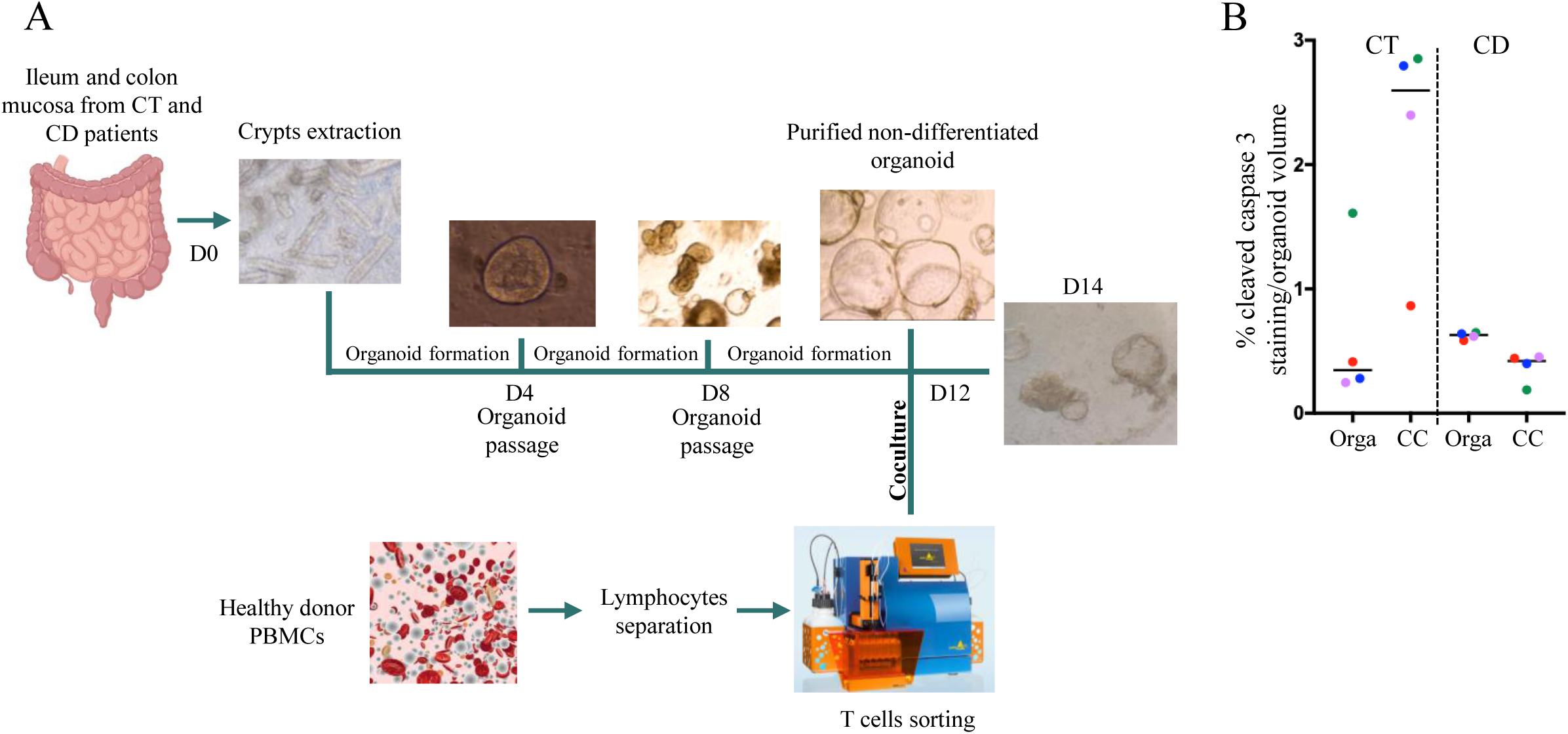
Allogenic cocultures between ileal organoids from CD patients or non-inflammatory control patients and peripheral T lymphocytes from healthy donor (A) Experimental design: purified ileal organoids are cocultured with peripheral T lymphocytes from healthy donor during 2 days before evaluation of epithelial cell death by confocal microscopy (B) Organoid epithelial cell death assessed with the cleaved Caspase-3 fluorescence volume over DAPI volume CC: cocultures, CT: control patients, CD: Crohn’s disease patients. All experiments were made in GM. Each point corresponds to the median of the values obtained for organoids of a given patient. Healthy donors n=4, CT patients n=4, CD patients n=4

### Mucosal T lymphocytes cocultured with purified ileal organoids reveal effector-like transcriptomic signature at single cell level

While the mechanisms underlying these results remain to be elucidated, they point to T cells are the main reason for the increase autologous IEC death in CD. Thus, to go further on understanding the immune mediated response observed in our experiments, we performed single cell RNA sequencing (scRNA-seq) of autologous T cells, cocultured or not with organoids from control and CD patients. T cells were recovered from the extracellular matrix and sorted by cytometry before single cell sorting and library preparation.

UMAP clustering showed defined clusters, which were then annotated based on the human immune system database. (Fig7.A) We then generated a heatmap of specific biomarkers to check the cell subtypes. As expected, we observed exclusively T cell subsets in our dataset: effector memory CD4 and CD8 T cells (Tem), central memory CD4 and CD8 T cells (Tcm), and two smaller groups of NK cells/γδ T cells and cycling Tem. (Supp2.A) The proportions of each cell subtypes were assessed in our samples one by one, and then compared between conditions: ML alone and ML cocultured, as well as between control and CD patients, with no major differences observed. (Supp2.B,C,D)

**Fig 7:**
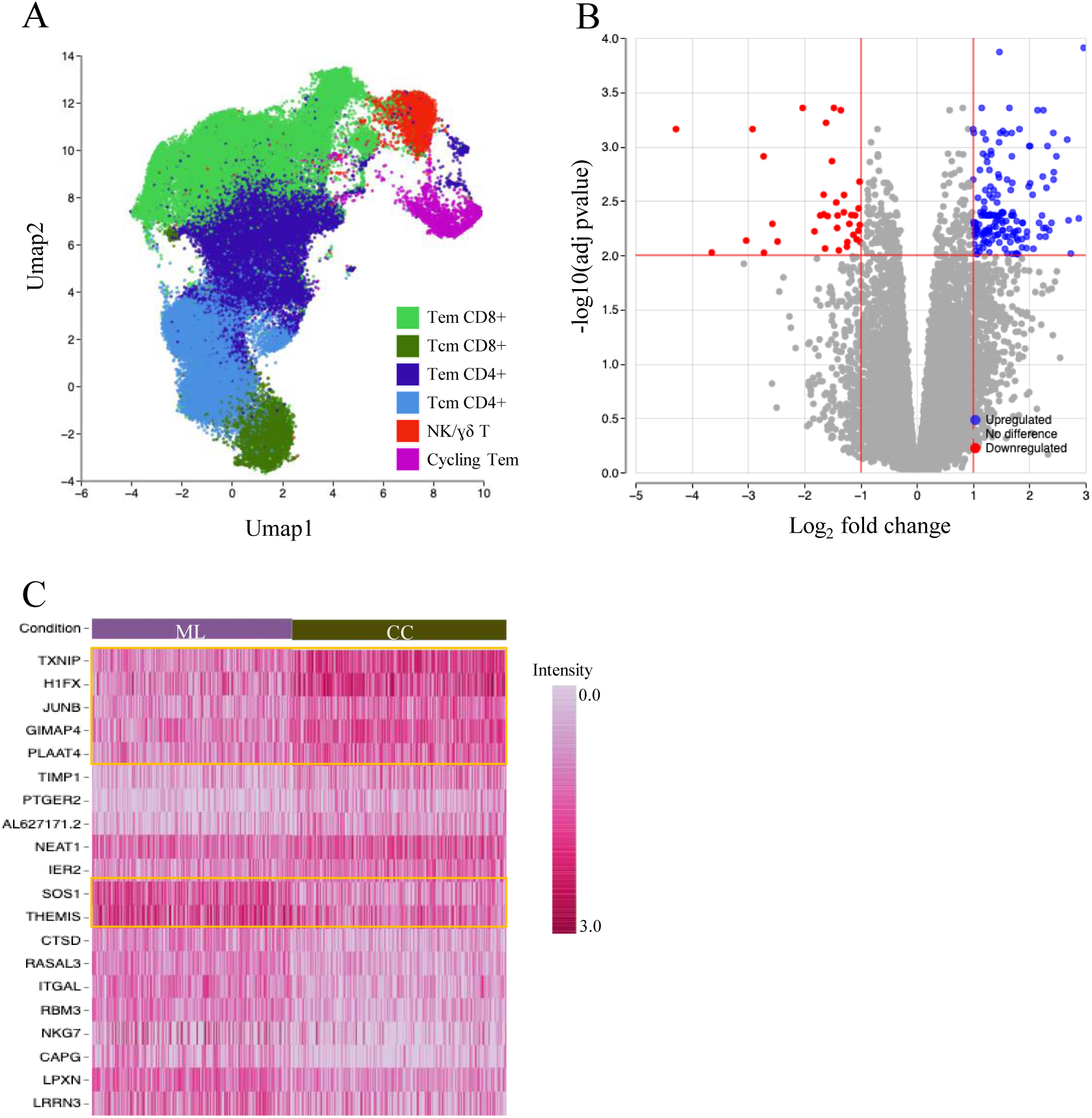
Autologous mucosal lymphocytes cocultured with ileal organoids present a particular gene signature. (A) Immune cell subtypes. UMAP of 66,349 CD5+ cells from control (n=4) and CD patients (n=4), colored by clustering and annotated post hoc (B) Volcano plot shows significantly up and downregulated genes in cocultured T cells compared to T cells alone (C) Heatmap shows cluster biomarkers that best characterize the group of ML alone and cocultured MLs Tem: effectors memory T cells, Tcm: central memory T cells, ML: mucosal lymphocytes, CC: cocultures, Orga: organoids alone. CT patients n=4, CD patients n=4

To next assessed the coculture effect, we compared the genes significantly up or down regulated in cocultured T cells compared to T cells alone (all patients and subtypes mixed). 163 up-regulated and 37 down-regulated genes were defined. (Supp3) Many of these genes were implicated in T cells regulatory mechanisms (such as GIMAP4, PLAAT4, DUSP1, ICAM4, GADD45B…) and metabolism (TXNIP, ABCG1, ABCA1, APOA1, THEMIS…). (Fig7.B,C) These results showed that T cells in our cocultures responded to the epithelial environment inducing pathways downstream of the TCR response.

We then tested if there were major differences between T cells from CD patients and from controls. Comparison of T cells alone from CD patients to T cells alone from controls showed no differentially expressed genes. (Supp4.A) The same way, the comparison between cocultured T cells from our two types of patients showed also no significant differentially expressed genes. (Supp4.B) Then, we compared all cells mixed from CD patients to those from controls: few differentially expressed genes were observed in T cells from CD patients (U2AF1L5, CCDC163, RHOD, TCL6…). (Supp4.C)

We then focused on the comparison between subtypes from CD and control patients and saw that 15 genes were downregulated in cocultured Tcm CD8+ from CD patients compared to the same group from controls. (Supp4.D) These results suggested that, overall, T cells from CD patients and controls were similar, but that differences can be detected by focusing on particular subpopulations, which could therefore be responsible for the differences observed in our cocultures.

We then asked if particular subpopulations could be identified in CD cocultures compared to control conditions that could explain what we observed in our experiments. Interestingly, when comparing the group of effector memory CD8 T cells (Tem CD8+) in coculture to the same population in control conditions, differential expression analysis showed 20 significantly up-regulated genes and 12 down-regulated genes. (Fig8.A,B) Among these genes, some up-regulated genes were related to T cell activation and Th1 responses like DUSP1, ATP2B1, GADD45B, JUN and PLAAT4 and some down-regulated were related to anti-inflammatory properties like ASB2, GPR68, PTPRJ, IL9R and AHRR. (Fig8.C) This Tem CD8+ population activated in cocultures presented thus an interesting pro-inflammatory profile, which should be implicated in the cytotoxicity and the proliferative defect observed in our experiments.

**Fig 8:**
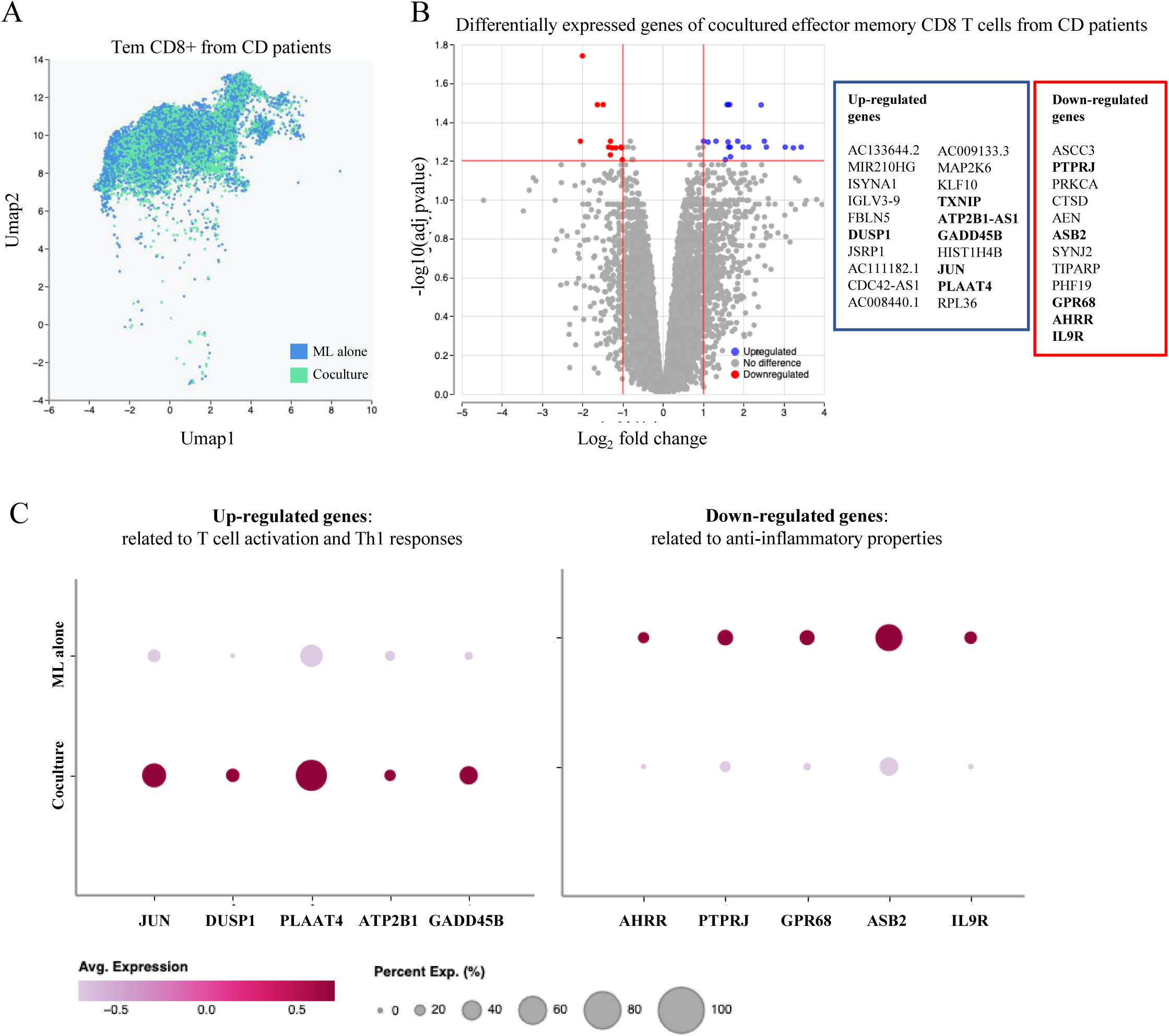
CD8 effector memory T cells from CD patients seem to be activated in coculture. (A) UMAP of Tem CD8+ cells from CD patients (n=4), colored by condition (alone or cocultured) (B) Volcano plot shows the genes differentially expressed in CD8 Tem cocultured from CD patients (C) Dot plots showing 5 up-regulated genes related to t cell activation and Th1 responses and 5 down-regulated genes related to anti-inflammatory properties ML: mucosal lymphocytes, Tem: effector memory T cells

## Discussion

Here, we highlighted the crosstalk between mucosal T lymphocytes and IECs that differentially impact epithelial maturation and survival at homeostasis and in inflammatory context. This autologous immune organoid cocultures model recapitulates the capacity of mucosal T lymphocytes to support epithelial differentiation while preserving ISCs pool in homeostatic context. More importantly, we demonstrated that, in the inflammatory context of Crohn’s disease, mucosal T lymphocytes acted aggressively against IECs. Epithelial cell death was observed, accompanied by a loss of organoids proliferative capacity and morphological changes. While control organoids were budding, those from CD patients in coculture had a decreased size and lost their budding structures.

We demonstrated that mucosal T cells support IECs differentiation towards secretory lineages at homeostasis, while preserving the ISC pool. But in the inflammatory context of CD, we observed an increased T cells-mediated IECs differentiation in parallel to a significant size decrease and a loss of budding. Since the total number of cells is decreased in coculture, two hypotheses were put forward to explain this observation: increased differentiation or decreased proliferative capacity of the organoids. This second hypothesis was tested, and we demonstrated a decrease of the Ki67+ proliferative cells, which were constant in control cocultures.

Although we were unable to demonstrate it directly, our results suggested a link between the epithelial death observed in coculture and the loss of proliferative capacity. Indeed, differentiated secretory cells were still present in organoids, probably resulting from early differentiation, while Ki67 proliferation marker was greatly reduced. ISCs and the resulting transit-amplifying cells are the proliferative cells expressing Ki67. A decrease in this marker suggested a loss of these populations, and more particularly of the ISCs pool.^25^ Moreover, the great capacity of intestine to repair after injury is well known. After the apoptosis phase to remove damaged cells, an active proliferation state follows to restore epithelium integrity. A possible revert process has been shown in differentiated IECs and +4 quiescent stem cells to replenish the ISCs pool. In this situation organoids lost their buds gradually to acquire a hyperproliferative phenotype allowing regeneration.^26^ However, this was not observed in our experiment, suggesting the killing of ISCs. In our model, even if this process of regeneration was initiated, it could not succeed if T cells targeted ISCs.

We wondered how much of these defects resulted from intrinsic modification in IECs from CD patients. Thus, we assessed their response to allogenic peripheral T cells from healthy donors. Surprisingly, no allogenic response was observed in our experiments with CD derived organoids, while control organoids were attacked. These results suggest that not only were they not more sensitive, but that they even had potentially inhibitory mechanisms to counter allogeneic responses, while still being susceptible to autologous mucosal T cells cytotoxicity. This then suggests that the increase in cytotoxic potential of T cells from CD patients was the main reason for the impaired lymphoepithelial interactions. Single cell RNA analysis of IECs from CD patients in literature suggest long-lasting defects in this cell type^27,28^, but to verify our hypothesis it would be interesting to perform transcriptomic analysis on our cocultured organoids. This would also potentially reveal pathways involved in lymphoepithelial interactions.

T cell lineage diversity in CD patient’s mucosa has been shown by single cell analysis and differences are observed with controls.^29^ But what interested us was to find out which particular population(s) extracted from the mucosa could explain the cytotoxic phenomenon observed. Then, deciphering the transcriptomic changes at single cell levels in our experiments, we were able first to confirm that T cells were indeed stimulated by the epithelial environment. But more interestingly, with a particular focus on the subtypes composing our cocultured T cell population, we highlighted the activation of effector memory CD8+ T cells in CD patients presenting a pro-inflammatory Th1 profile. This particular population was not highlighted in cocultures from control patients, suggesting its central role in the immune mediated cytotoxicity observed in our experiments in inflammatory context. Now we have demonstrated the cytotoxic potential of mucosal T lymphocytes all together, it could be interesting to perform cocultures with sorted subsets. This would either show that a single population is responsible for the cytotoxicity observed, or that dialogue is necessary.

Autologous T cell cytotoxicity in CD is sustained despite organoid passages, as purified organoids are free of all other cell types or bacterial remains that could have influenced our previous observations.^30^ Moreover, providing an inflammatory signal by stimulating purified organoids with LPS did not increase their sensitivity to T cells. (Supp5) The nature of the interactions has yet to be fully dissected, although we showed the involvement of NKG2D and CD103 in lymphoepithelial interactions and indirect role of cytokines.^22^ The question of the potential antigen(s) involved remains. The direct involvement of the TCR is still in question as genes known to be downstream of the receptor were not significantly modulated. Hence, the deleterious interactions could very well be TCR-independent. The use of MHC-blocking antibodies in our cocultures could provide us with answers.^31^ TCR sequencing of mucosal T cells in coculture could also bring us responses regarding the nature of lymphoepithelial interactions, antigens involved and potential clonal expansions.

## Conclusion

Here, we demonstrate the capacity of mucosal T lymphocytes from CD patients to impair the epithelial integrity of organoids in an autologous coculture model. Our data suggest that pro-inflammatory CD8+ effector Trm are drivers of the cytotoxicity against ISCs. This model can be used to investigate tissue-resident immune responses at homeostasis and to test immune based T cell targeting therapies in the context of CD.

### Material and Methods

### Introduction of cohorts

#### The POP-REMIND cohort

The POP-REMIND cohort included patients since September 2010 based on the following criteria: be over 18 years old, having an ileal or ileocolonic CD and indication of CD-related intestinal surgery (ileocolonic resection). Six months after surgery, all patients had a colonoscopy to assess the endoscopic recurrence according to the Rutgeerts score. Ileal and colon mucosa samples were collected from the resection piece for experimentations. (Table 1)

**Table 1:**
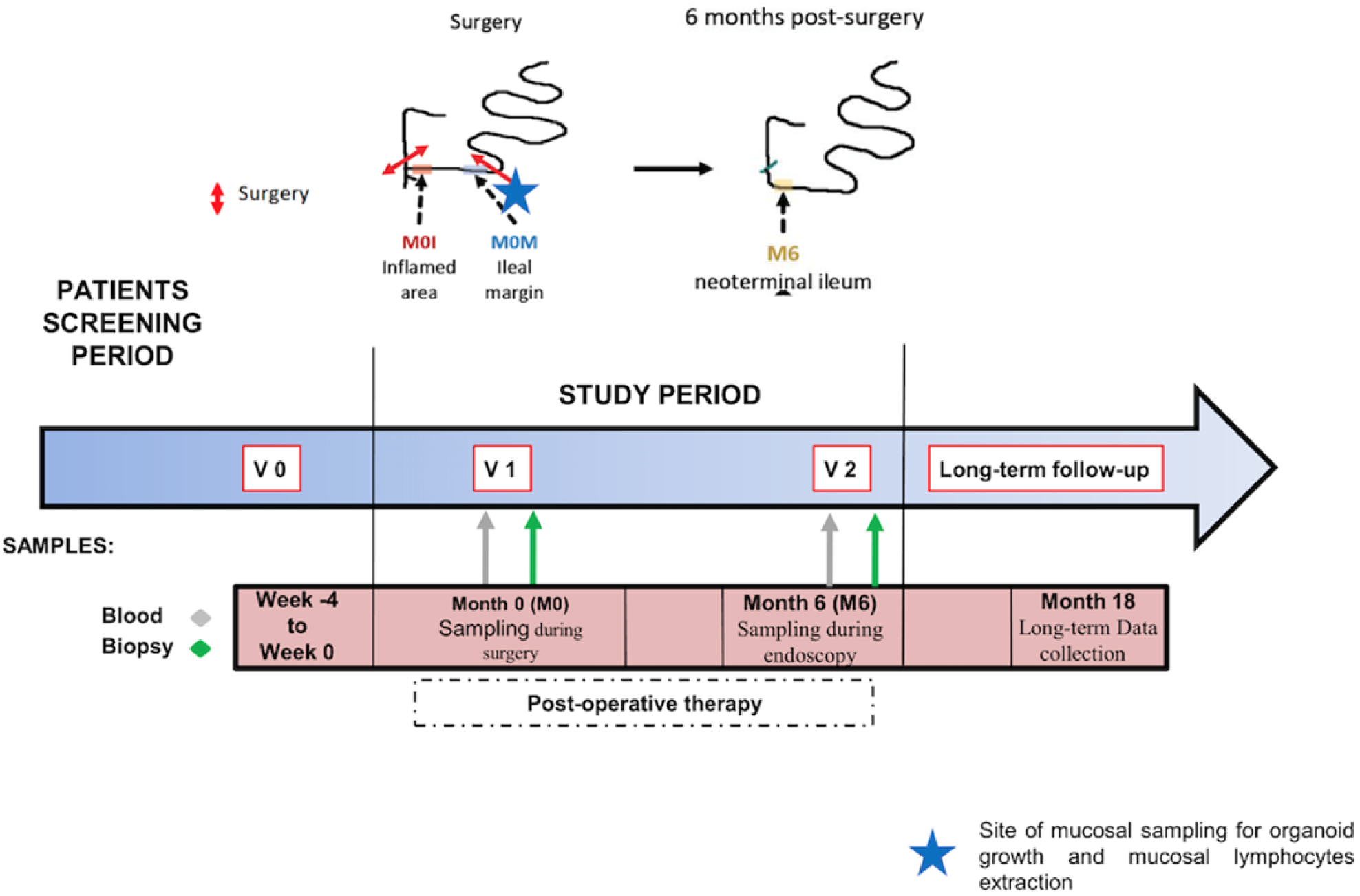
Study design of POP-REMIND cohort (from M. Bezault)

#### The IMCO cohort

The IMCO cohort included patients prospectively since February 2017 who underwent resection for colon cancer at the Saint-Louis Hospital. For patients with right colon cancer who underwent colectomy, ileum mucosa was then available for experimentations, in addition to the colon mucosa.

### Autologous coculture between intestinal organoids and mucosal lymphocytes from patients

#### Mucosal lymphocytes isolation and culture

Samples of ileal mucosa were retrieved from the resection pieces of patients included in the cohorts. After several washes with PBS, to avoid contamination, the samples cut in small pieces were incubated under agitation 20 minutes at room temperature in PBS containing fungizone (Gibco, 15290), gentamicin (Lonza, 17-519Z) and normocin (Invivogen, ant-nr-1). After washing, the fragments were enzymatically and mechanically digested with collagenase IV (Sigma-Aldrich) and DNase (Roche) in complete RPMI medium at 37°C under agitation during 4×7 minutes separated by vigorous shake for 1 minute. Collagenase action was stopped by adding 10mL of cold Fetal Bovine Serum (FBS). The supernatants were filtered and centrifugated at 1500rpm for 5 minutes. The cell pellet was layered on a density gradient by Ficoll separation method. After centrifugation, the immune cells were collected from the interface, washed with PBS, and resuspended in complete RPMI to be counted on a Kova slide. The cells were then cultured with IL15 and IL7 (Milteyni) at the concentration of 10ng/mL in a 96 round bottom plate with 200 000 cells per well to maintain them alive until their use.

#### Flow cytometry analysis of isolated mucosal lymphocytes

A sample of resting T lymphocytes was stained with the following antibody panel and analyzed by flow cytometry at the day of the coculture (from BD Biosciences and Miltenyi). (Table 2) The cells were also co-stained with DAPI, all diluted in optimal concentration in FACS Buffer (Miltenyi Biotec). The cell suspension was analyzed with an Attune NxT flow cytometer (Life Technologies) and the data obtained were analyzed with FlowJo software (Tree Star).

**Table 2:**
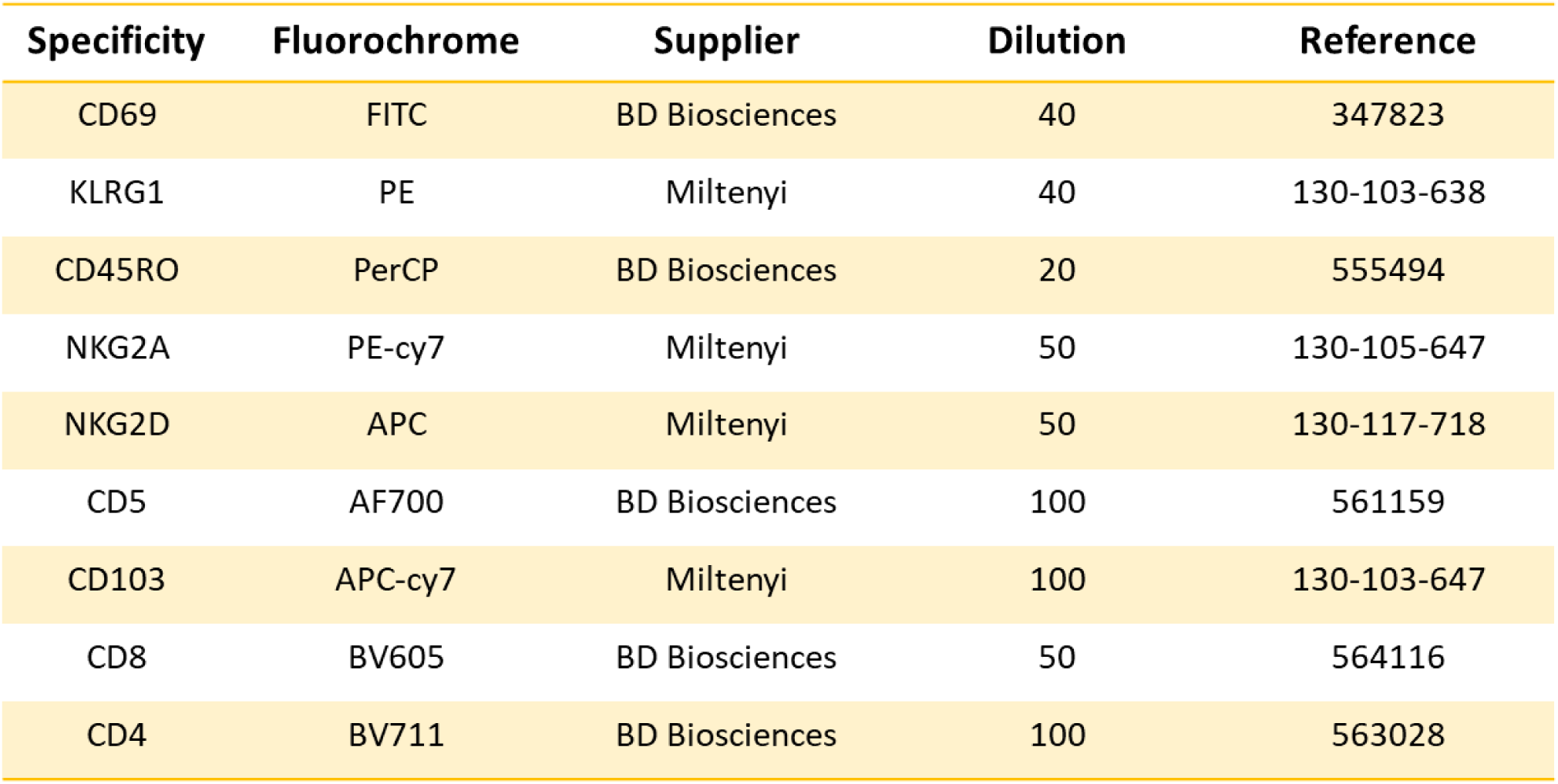
Mucosal lymphocytes Flow cytometry panel applied after extraction and at coculture time.

#### Intestinal crypts extraction and organoid formation

Samples of ileal and colon mucosa were cut into small fragments. The fragments were incubated in PBS containing fungizone (Gibco, 15290), gentamicin (Lonza, 17-519Z) and normocin (Invivogen, ant-nr-1) 25 minutes under agitation at room temperature. After washing, they were incubated 7 minutes with mucolytic agents dithiothreitol (DTT, Sigma-Aldrich) at the concentration of 0.5mM at room temperature under agitation to remove the mucus. After washing, they were incubated 45 minutes with Ethylene Diamine Tetraacetic Acid (EDTA 0.5M, Invitrogen) at the concentration of 8mM on ice under agitation to fragilize the crypts attachment by chelation. After washing 25mL of cold PBS were added and the tube was vigorously shaked. The supernatant was recovered in 50mL Falcon tube containing 5mL of cold FBS. This operation was repeated at least 2 times. The different fractions obtained were centrifugated 3 minutes at 150rcf at 4°C. The cell pellet was resuspended in 1mL of complete medium RPMI and number and quality of crypts obtained were evaluated under microscope on a petri dish. The fraction containing the most well-formed crypts and the least mucus was centrifugated 3 minutes at 150 rcf. On ice, the cell pellet was resuspended in Matrigel matrix (Corning) to form 20µL drops containing around 100 crypts in a 24 round bottom plate (ThermoFisher Scientific). The plate was incubated 20 minutes at 37°C to polymerize the Matrigel. 200µL of Intesticult Organoid Growth Medium (OGM, Stemcell Technologies) supplemented with 5mM of anoikis inhibitor Y27632 and Penicillin streptomycin (Gibco) was added in each well. The media was changed at day 1 then each 3 days with OGM. At this point, mixed (non-passaged) organoids were obtained.

Once the mixed organoids had sufficiently grown (after five days approximately), they were “passaged”. Medium was removed and 200uL of Gentle cell dissociation reagent were added to digest Matrigel drops during 5 minutes at room temperature. The enzyme action was stopped by adding 1mL of cold RPMI medium in each well. Organoids were collected and centrifugated 3 minutes at 300rcf. Supernatant was eliminated and 1mL of cold RPMI medium was added. With a 1mL pipet, organoids were mechanically broken by flushing them to obtain small clusters of intestinal epithelial cells. The suspension was then filtered with a 100uM pore filter to eliminate what was not disrupted. Clusters of IECs were put back in culture in 20uL drop of Matrigel matrix with around 100-150 clusters per drop in a 24 round bottom plate (ThermoFisher Scientific). The plate was placed at 37°C for Matrigel polymerization for 20 minutes. 200µL of Intesticult OGM supplemented with 5mM of anoikis inhibitor Y27632 and Penicillin streptomycin was added in each well. Passages were repeated until purified organoids composed mainly of ISCs were obtained, characterized by their giant bubble shape.

Autologous coculture between mucosal lymphocytes and organoids

All the components were conserved on ice during the manipulation. MLs were recovered from the 96 well plate, washed, and counted on a Kova slide. The medium of organoid wells was removed, and each drop was incubated 5 minutes at room temperature with 200uL of Gentle cell dissociation (Stemcell Technologies) to digest the Matrigel. Digested drops were recovered and washed in complete RPMI to count the total organoid number under microscope on a Petri dish. For the coculture MLs and organoids were mixed with a ratio of one organoid for 10 000 MLs. For cocultures dedicated to microscopy, flattened drops of 10uL of Matrigel containing 10 organoids for 100 000 MLs were made into IBIDI u-Slide 8 wells (Biovalley). For cocultures dedicated to sorting and FACS, drops of 20uL Matrigel each containing 20 organoids for 200 000 MLs were made into 48 bottom plate. IBIDI and plates were then placed at 37°C for Matrigel polymerization for 20 minutes. The wells were then filled either with Intesticult OGM or Organoid Differentiation Medium Intesticult (ODM, Stemcell Technologies) to induce differentiation by the medium. The plates were then incubated at 37°C for 5 days.

At this point the use of Differentiation medium was questioned as it tended to erase differences between our conditions and influence organoid size. This medium is notably composed of DAPT, an inhibitor of the ỿ-secretase essential in Notch pathway activation. Its inhibition is known to lead to a decrease of proliferation, a loss of ISCs, an increased cell apoptosis and an early differentiation of progenitors into secretory cells. We thus decided at some point of our study to focus only on results with Growth medium.

#### Confocal microscopy: staining, acquisition and ICY analysis

The supernatants were removed from IBIDI cocultures and stored at −80°C. To fix the coculture, the wells were washed with PBS. 200uL of Paraformaldehyde (PFA, Sigma-Aldrich) 4% were added and removed after 20 minutes of incubation at 37°C. The cells were then permeabilized by adding 200uL of permeabilization solution composed of Triton X-100 (Sigma-Aldrich) 0.1%, incubated at room temperature for 45 minutes. 200uL of the blocking solution composed of Bovine Serum Albumin (BSA, Sigma-Aldrich) 0,1% was added in each well and incubated for 2 hours at room temperature. Then 170uL of primary staining antibody mix made with BSA blocking solution 0,1% were added and incubated overnight at 4°C. Then 170uL of the secondary staining antibody mix and DAPI made with BSA blocking solution 0,1% were added and incubated 2 hours in dark at room temperature. (Table 3) Finally, 170ul of the transparization solution (Rapiclear 1.47 SunJin Lab) were added. IBIDIs were conserved at 4°C in dark until microscope acquisition.

**Table 3:** Primary and secondary antibody mix used for coculture staining.

| Primary antibodies | Secondary antibodies |
| --- | --- |
| Ms mAb to Muc2 (ab11197, Abcam) | Goat anti-mouse Af488 (ab150113, Abcam) |
| Rb mAb to Caspase 3 (ab4051, Abcam) | Goat anti-rabbit Af647 (ab150083, Abcam) |
| Rat mAb to CD3 (ab11089, Abcam) | Goat anti-rat Af555 (ab150158, Abcam) |
| Rb mAb to Lysozyme (ab108508, Abcam) |  |
| Ms mAb to Ki67 (ab245113, Abcam) |  |
| Rb mAb to Ki67 (ab92742, Abcam) |  |

Confocal spinning disk microscope (Inverted Eclipse Ti, Nikon; Confocal system: CSUX1-A1N, Yokogawa) was used to image the experiments. Icy software was used to analyze the data collected. Six parameters were quantified: the organoid size (DAPI volume), the epithelial cell death percentage (cleaved caspase 3 volume over the DAPI volume of organoid), the number of CD3+ cells interactions (counting of the number of CD3+ cells in contact with organoid, or close, over the DAPI volume of organoid), the percentage of Goblet cells number (counting of the number of MUC2 positive cells over the DAPI volume of organoid), the percentage of organoids with lysozyme positive staining and the percentage of proliferative capacity (Ki67 volume over the DAPI volume of organoid). Within the same experiment, for DAPI, cleaved caspase 3 staining and Ki67 approximately the same thresholds were used for all organoids to obtain the most specific staining possible (Threshold tool). Fluorescent area was extracted (Label extractor) and ROI statistics were processed to obtain the volume of staining. (Table 4) For each parameter of each patient in each condition the median value of all organoids was calculated.

**Table 4:**
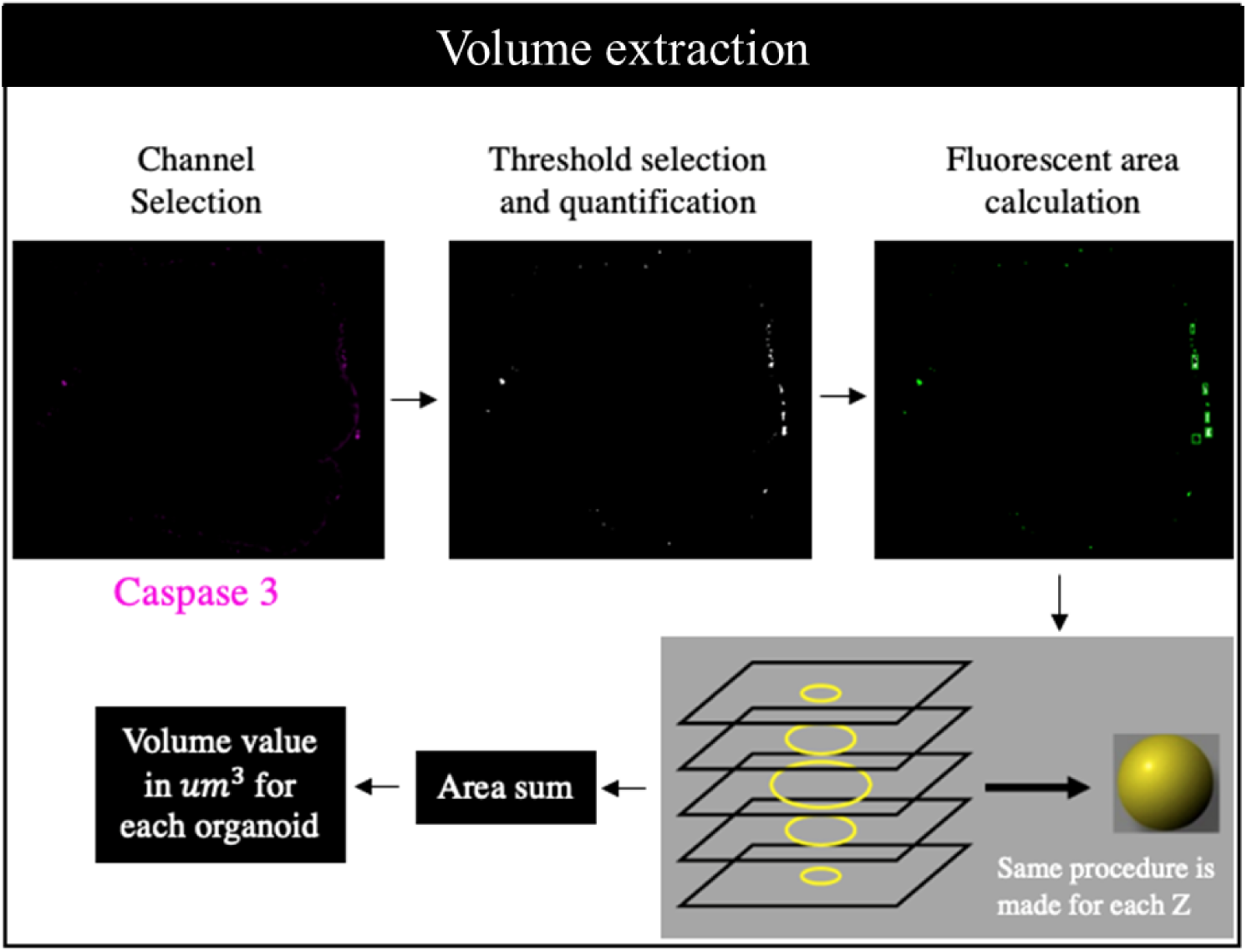
Calculation of cleaved caspase3 labelling volume with Icy software for all Z-stack of one organoid imaged by confocal microscopy.

#### Obtaining a cell suspension from cocultures for flow cytometry and transcriptomic analysis

Supernatants were removed from plates cocultures and stored at −80°C. All the components were conserved on ice during the manipulation. Drops of ML alone, organoids alone or cocultures with autologous T lymphocytes were incubated 5 minutes at room temperature in 200uL of Gentle cell dissociation reagent. The cells were recovered and washed in cold complete RPMI medium. After centrifugation at 300rcf 3 minutes at 4°C the pellet was resuspended in 1mL of cold complete medium and organoids were break mechanically by flushing them to obtain small clusters of cells. The suspension was centrifugated at 150rcf 3min at 4°C. After elimination of the supernatant, 500uL of Trypsin (Gibco) supplemented in Y were added. The cells were incubated 6 minutes at 37°C. The clusters were flushed to obtain a cell suspension. The trypsin action was stopped by adding 500uL of cold complete medium supplemented in Y. The suspension was then filtered with a 100uM pore filter to eliminate what was not disrupted. A little sample of the cell suspension was put aside to perform qRT-PCR of Ki67, LGR5 and Sox9 genes. The rest of the suspension was centrifugated 3 minutes at 1500rpm and 50uL of the fluorescent-labeled antibody panel was added.

For proliferation phenotyping on Attune NxT flow cytometer, anti-EpCAM and anti-CD45 staining were used. The cells were stained 20 minutes in dark at 4°C. Then they were stained with a viability marker for 10 minutes at 4°C. The cells were then fixed and permeabilized for 30 minutes at 4°C. An internal staining of Ki67 was then made for 30 minutes at 4°. The cells were finally stained with DAPI and processed on the flow cytometer. (Table 5)

**Table 5:** Flow cytometry panel used to sort T cells and IECs from coculture.

| Specificity | Fluorochrome | Supplier | Dilution | Reference |
| --- | --- | --- | --- | --- |
| CD45 | PE | BD Biosciences | 40 | 555483 |
| EpCAM | APC | Miltenyi | 50 | 130-111-000 |
| CD5 | AF700 | BD Biosciences | 100 | 561159 |
| CD8 | BV605 | BD Biosciences | 50 | 564116 |
| CD4 | BV711 | BD Biosciences | 100 | 563028 |

For the cell sorting on Aria III the cell suspension was stained 45 minutes in dark at 4°C with anti-EpCAM, anti-CD45 and anti-CD5 antibodies. After a wash the cells were stained with DAPI. The cells were gated on live DAPI negative cells to sort on one side the CD5 positive cells and on the other side the EpCam positive CD45 negative cells. (Table 6)

**Table 6:**
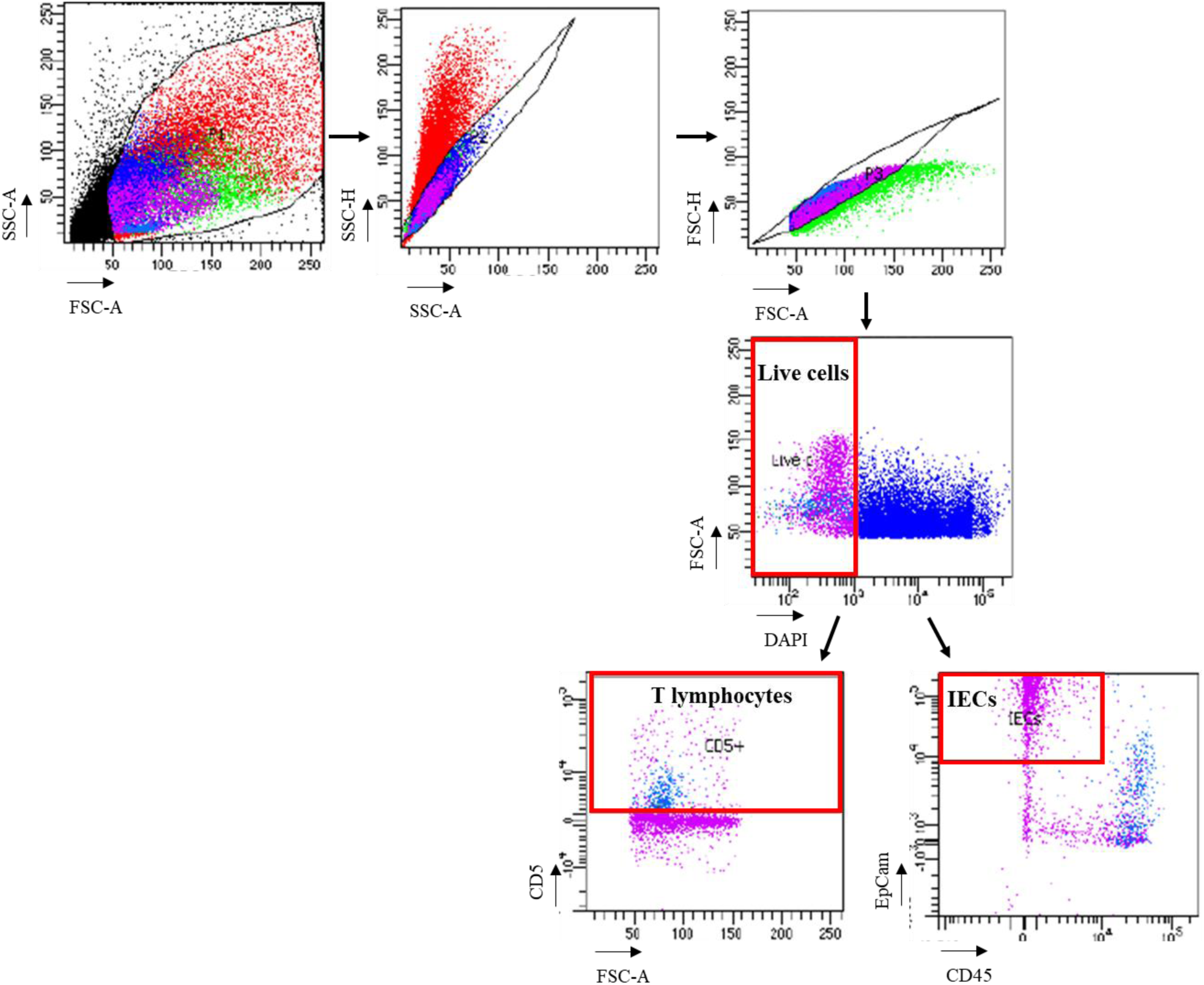
Gating strategy for mucosal T lymphocytes and IECs sorting by flow cytometry.

#### Protein analysis of coculture supernatants

Protein from frozen coculture supernatants were analyzed with the Luminex technology (Thermofisher). Two different kits (procartaplex and high sensitivity procartaplex) were used on coculture supernatants. (Table 7)

**Table 7:**
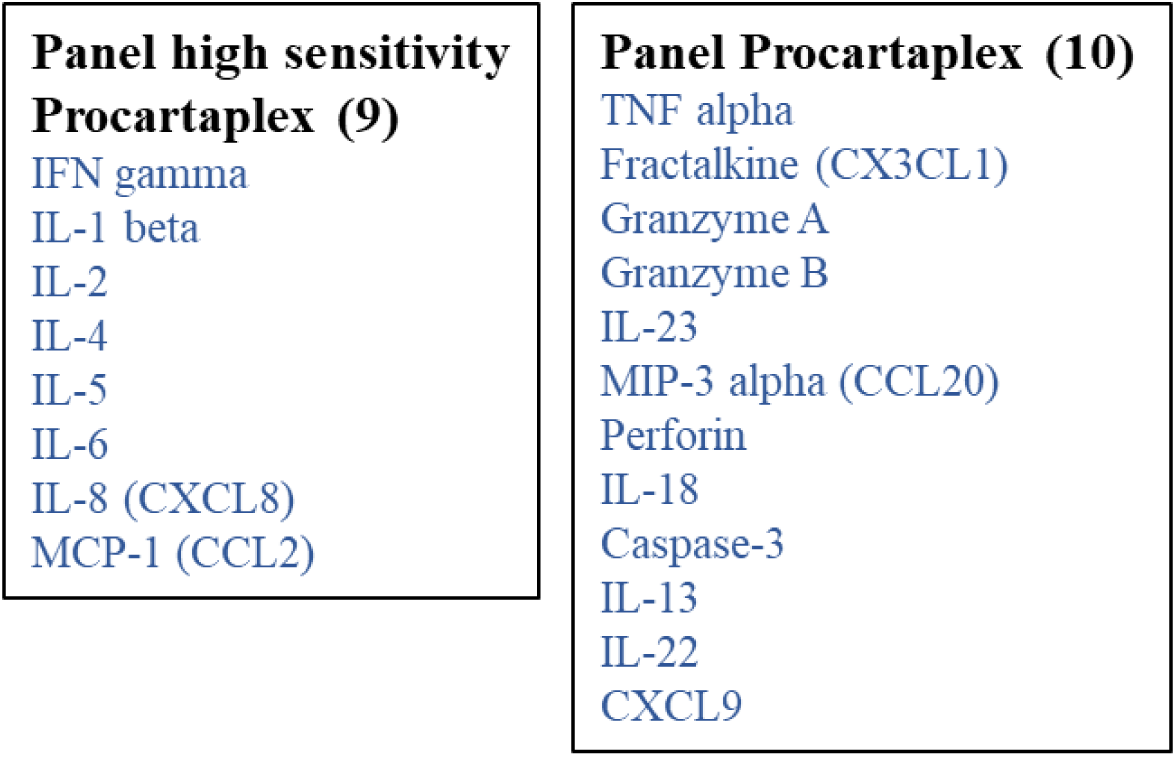
Cytokine panels for protein analysis of coculture supernatants.

#### Single cell TCR and RNA transcriptomic analysis of cocultured T cells

Single cell TCR-RNA sequencing experiments were performed with 10X Genomics technology on mucosal T lymphocytes. T cells were sorted from autologous coculture with organoids or from culture alone. Four cocultures of control patients and four cocultures of CD patients were performed. For each, 8000 cellules were targeted for sequencing. 10X genomics recommendation was to sort at least 50 000 cells. (Table 8)

**Table 8:**
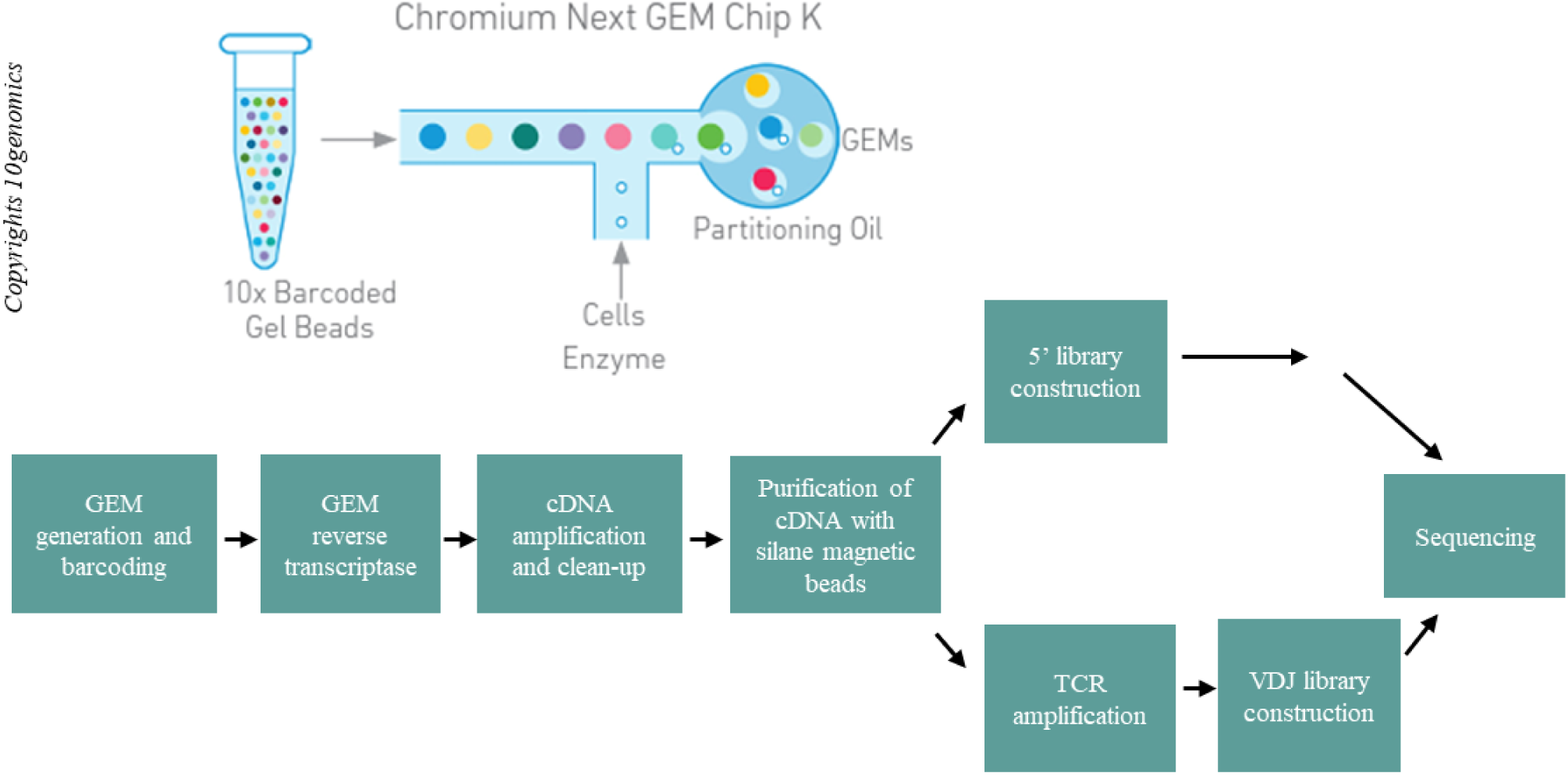
Workflow summary of single cell experiments.

The sorting was performed on FACS Aria III. After elimination of doublets, T cells were gated on live cells (DAPI negative), CD5 positive, EpCAM negative. Chromium controller was then used for the GEM generation and barcoding. The final libraries were sequenced on Illumina Novaseq, with 40 000 reads depth for the 5’ gene expression profile and 5000 reads depth for the VDJ.

### Statistical analysis (Prism, R)

Prism software was used to apply paired Wilcoxon or unpaired Mann-Whitney tests to analyze confocal microscopy, flow cytometry, Luminex, and qRT-PCR results on cocultures.

To analyze single-cell RNA sequencing data we used Cellenics software developed by Bioimage.

## Supplementary data

**Supp 1:**
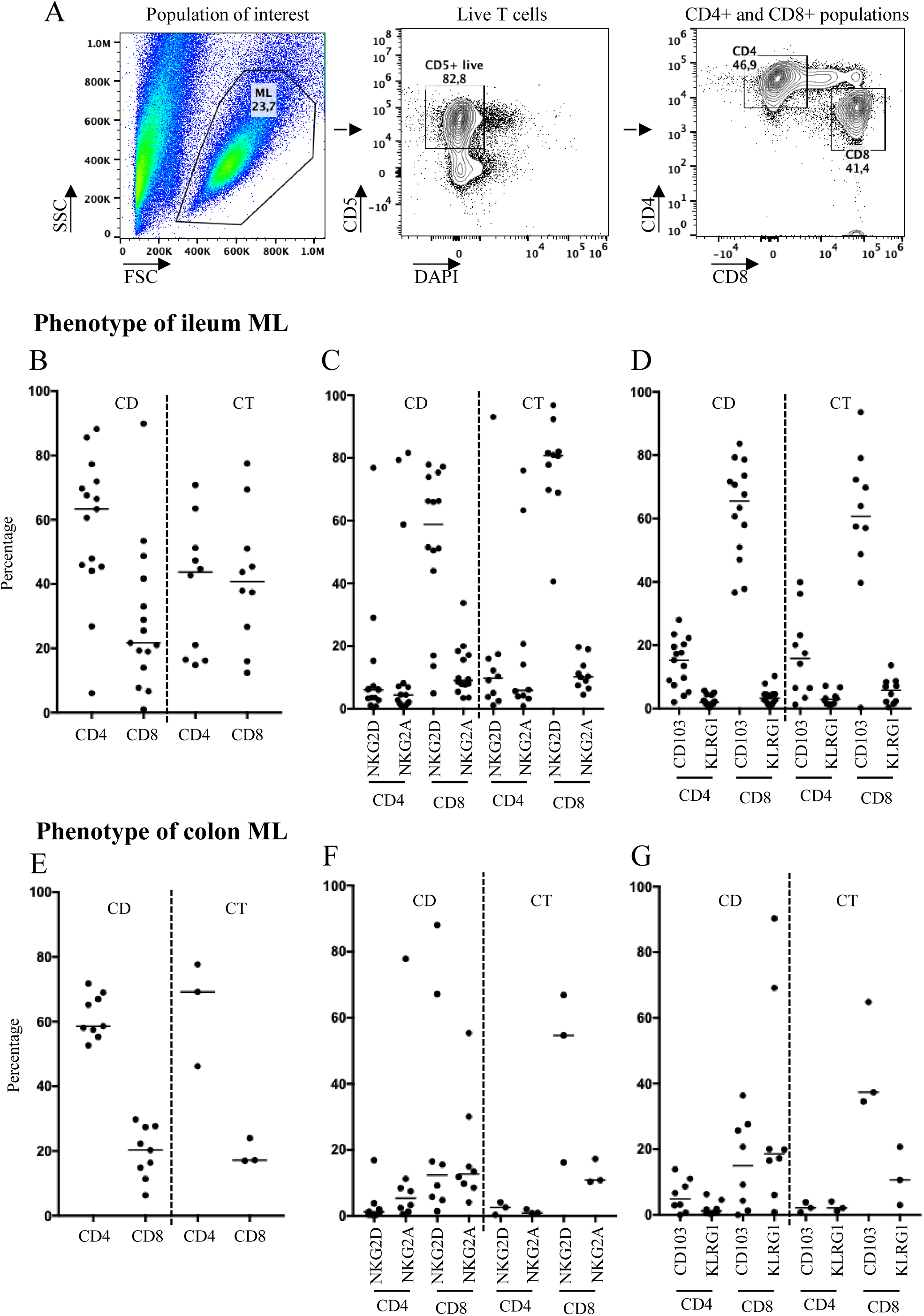
Ileal mucosal lymphocytes of CD patients are enriched in CD4+ T cells. (A) Gating strategy to assess viability of ML and their phenotype after long-term culture (B) Percentage of CD4+ and CD8+ cells in ileum ML (C) Percentage of NKG2D+ and NKG2A+ cells in ileum ML (D) Percentage of CD103+ and KLRGI+ cells in ileum ML (E) Percentage of CD4+ and CD8+ cells in colon ML (F) Percentage of NKG2D+ and NKG2A+ cells in colon ML (G) Percentage of CD103+ and KLRGI+ cells in colon ML

**Supp 2:**
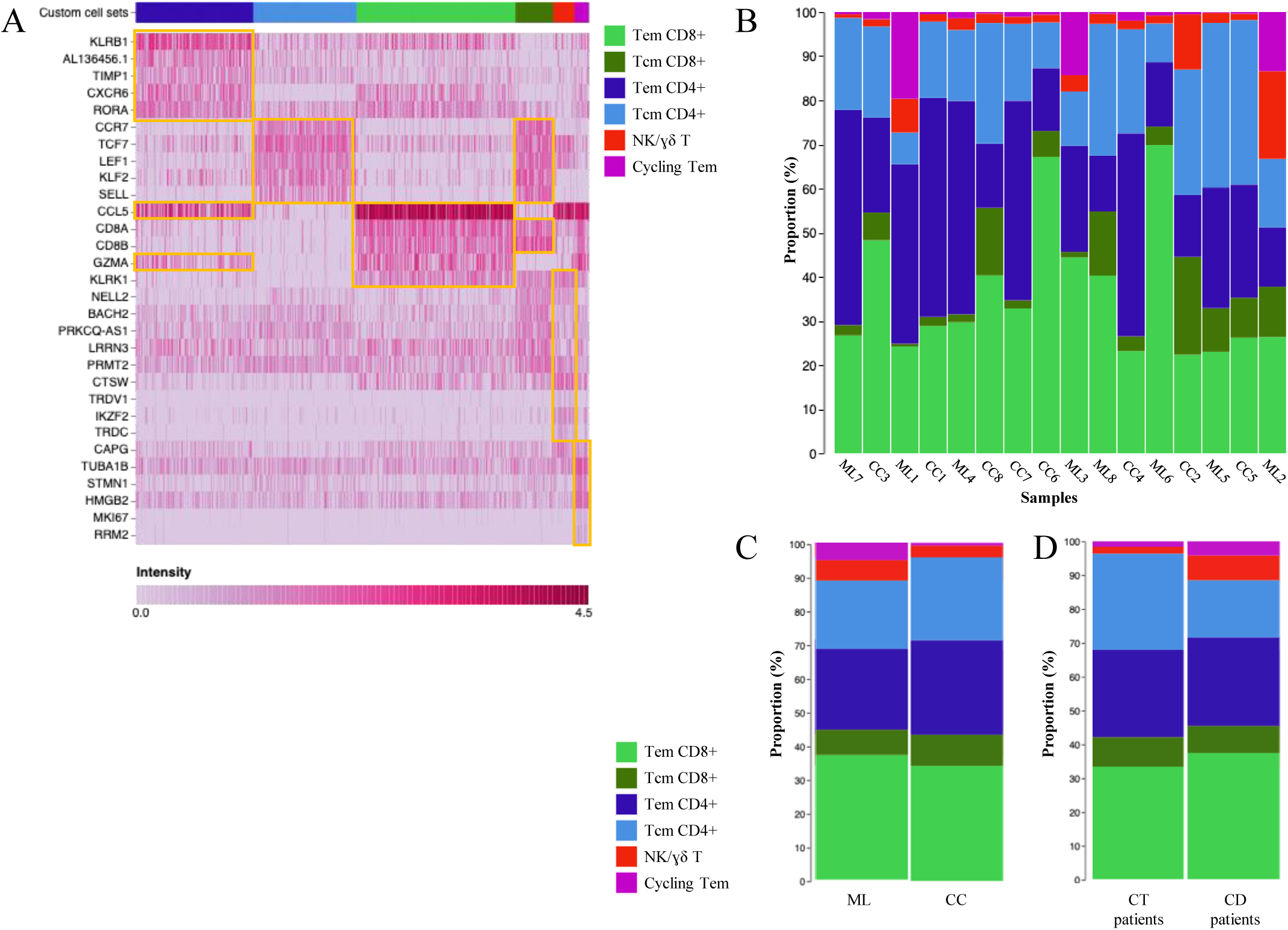
Single-cell population analysis. (A) Heatmap shows cluster biomarkers to identify cell subtypes (B) Proportion of cells from each identified cluster in each of the samples (C) Proportion of cells from each identified cluster for each of the two conditions (ML alone or cocultured) (D) Proportion of cells from each identified cluster for each of the two type of patients (control or CD patients) Tem: effectors memory T cells, Tcm: central memory T cells, ML: mucosal lymphocytes, CC: coculture, Orga: organoids alone. CT patients n=4, CD patients n=4

**Supp 3:**
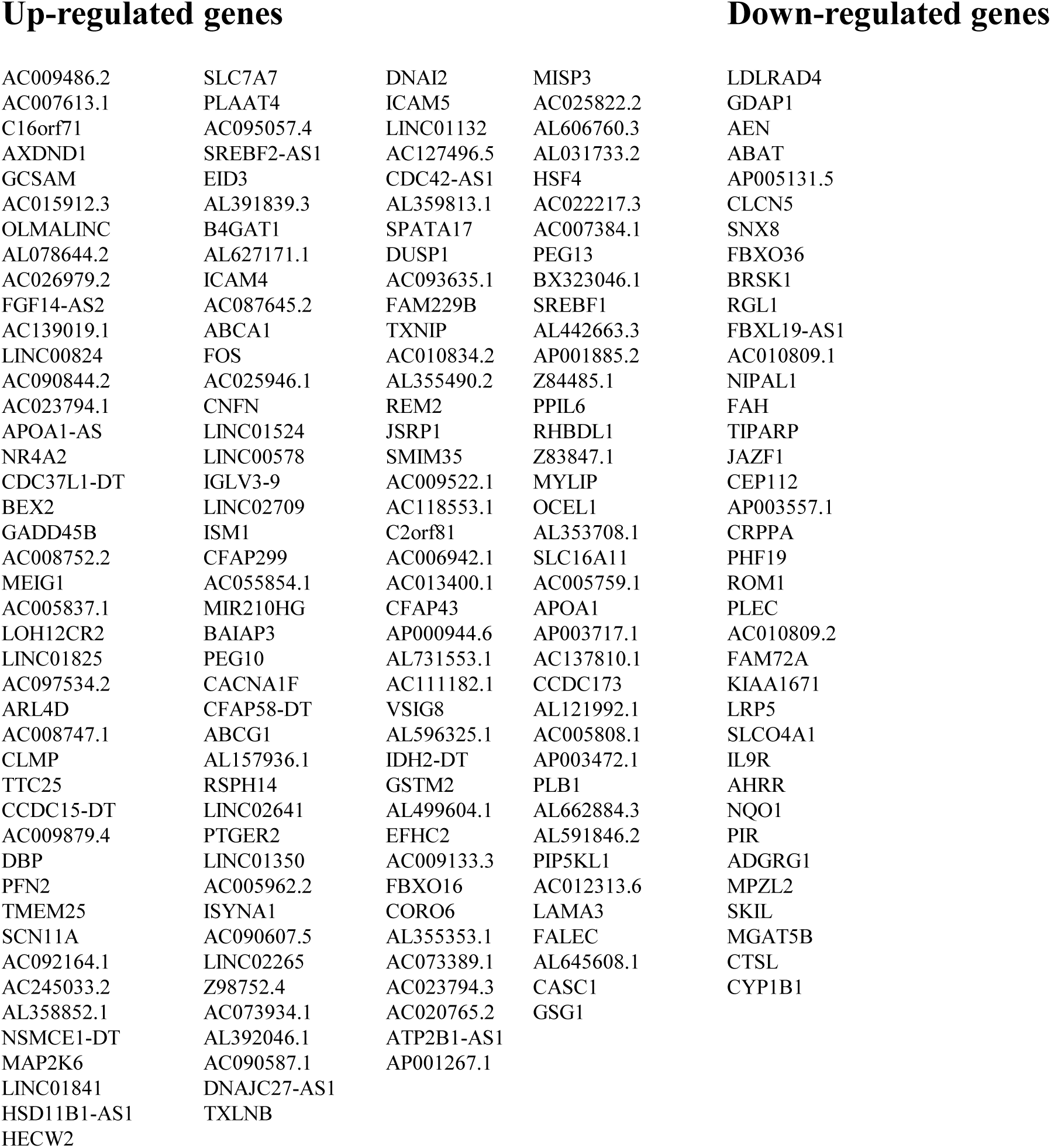
Significantly expressed genes (pvalue<0.01) in cocultured T cells compared to T cells alone.

**Supp 4:**
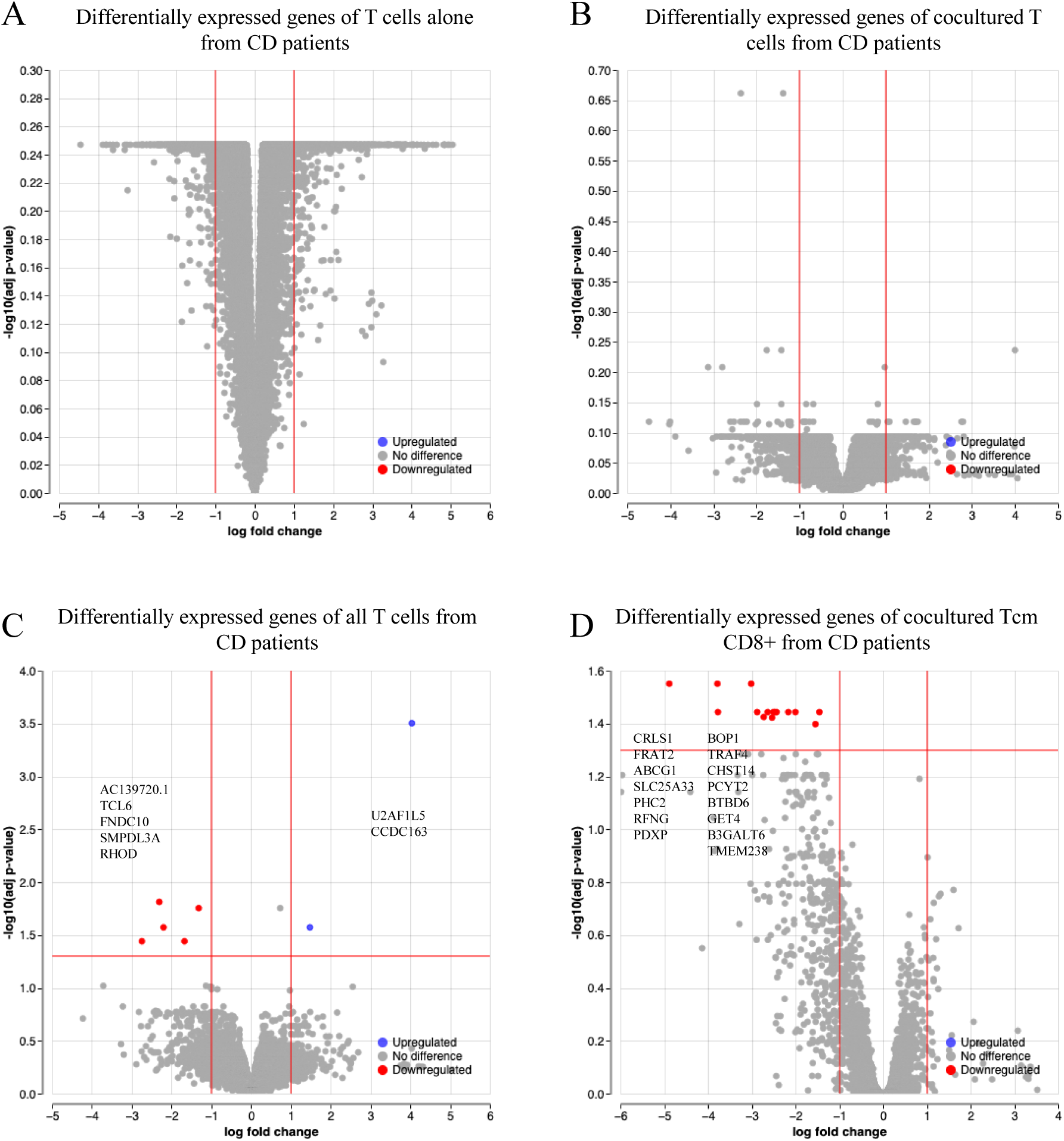
Autologous mucosal lymphocytes from CD patients present comparable gene signature to those from control patients. (A) Volcano plot shows no significant differentially expressed genes in T cells alone from CD patients compared to controls (B) Volcano plot shows no significant differentially expressed genes in cocultured T cells from CD patients compared to controls (C) Volcano plot shows significantly up and downregulated genes in all T cells from CD patients compared to controls (D) Volcano plot shows significantly downregulated genes in cocultured Tcm CD8+ from CD patients compared to controls

**Supp 5:**
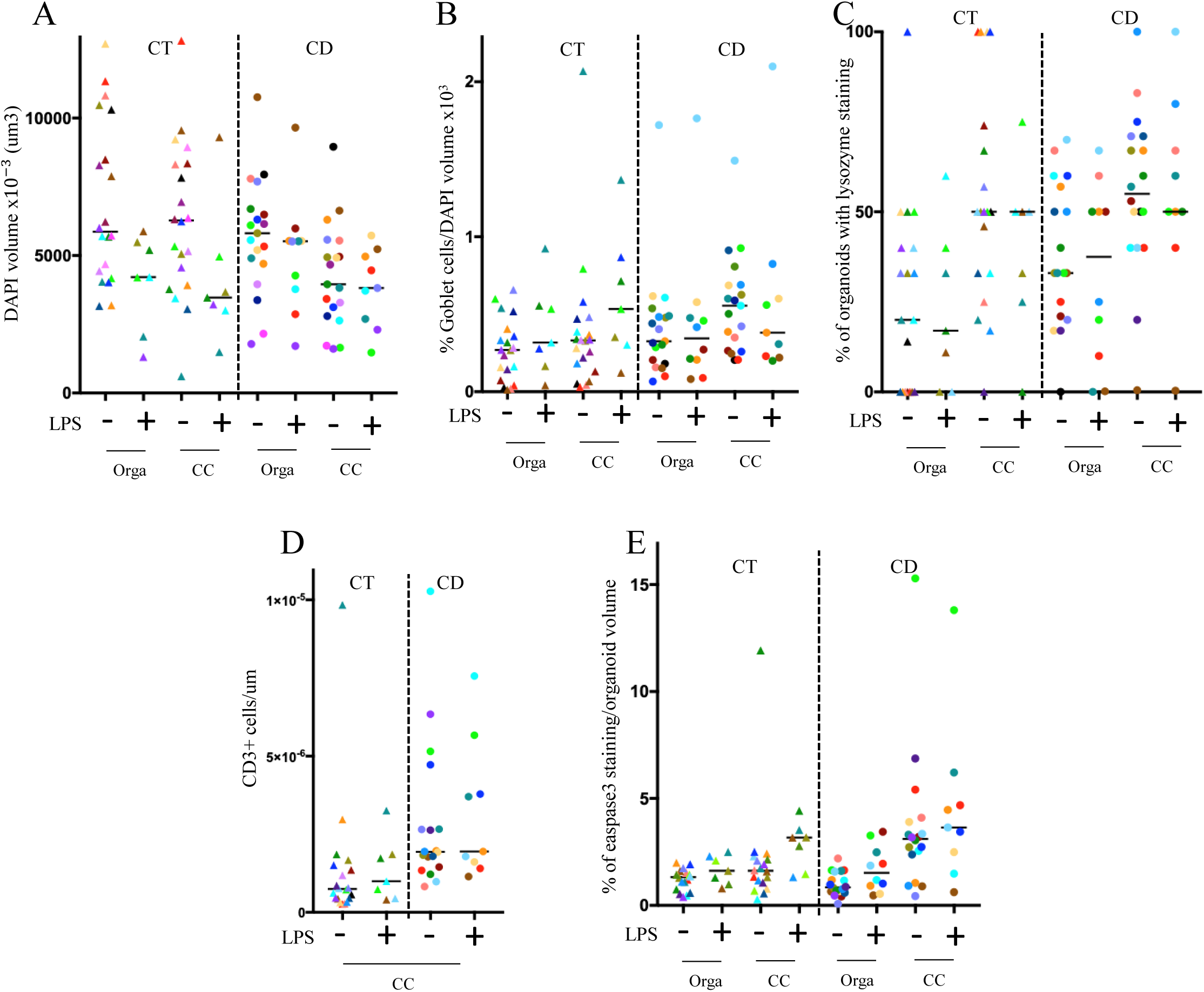
LPS does not increase CD patient organoids sensitivity after several passages. (A) Quantification of ileal organoids size through DAPI volume staining (B) Quantification of the percentage of Goblet cells over the DAPI volume and percentage of organoids with lysozyme (C) Organoid epithelial cell death assessed with cleaved Caspase-3 fluorescence area over DAPI volume (D) T cell infiltration assessed with the number of CD3+ T cells in contact over DAPI volume CT: organoids from control patients, CD: organoids from CD patients, CC: coculture, Orga: organoids alone All experiments were made in GM. Each point corresponds to the median of the values obtained for organoids of a given patient.

**Supp 6:**
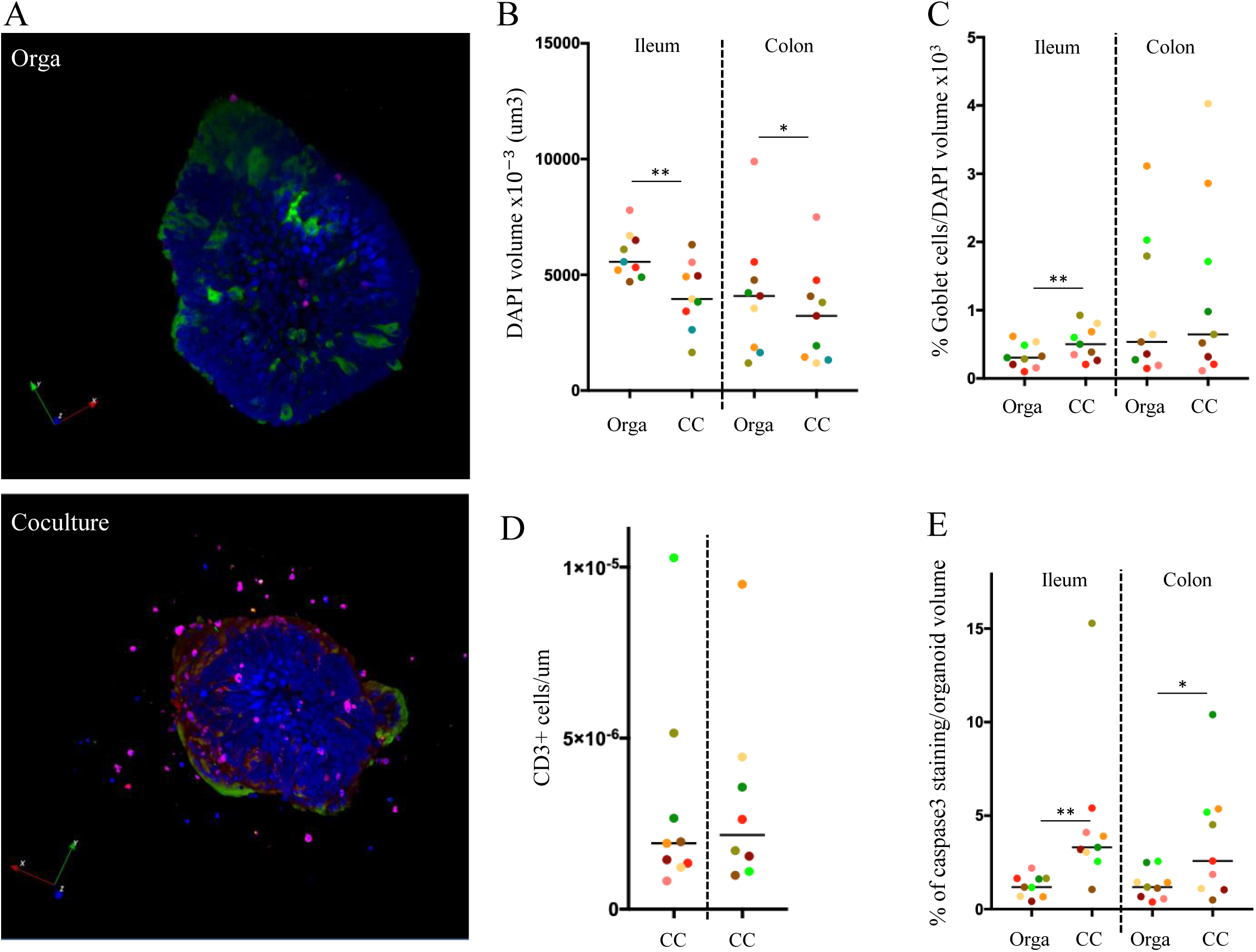
A similar immune mediated pattern is observed in colon organoids from CD patients. (A) Confocal microscopy (200X) of autologous colon coculture from CD patient alone or in coculture (DAPI: blue, Muc2: green, Caspase3: pink, CD3+ cells: red) (B) Quantification of ileal and colon organoids size from CD patients through DAPI volume staining (C) Quantification of the percentage of Goblet cells over the DAPI volume (D) T cell infiltration assessed with the number of CD3+ T cells in contact over DAPI volume (E) Organoid epithelial cell death assessed with the cleaved Caspase-3 fluorescence volume over DAPI volume All experiments were made in GM. CC: coculture. Each point corresponds to the median of the values obtained for organoids of a given patient.

**Supp 7:**
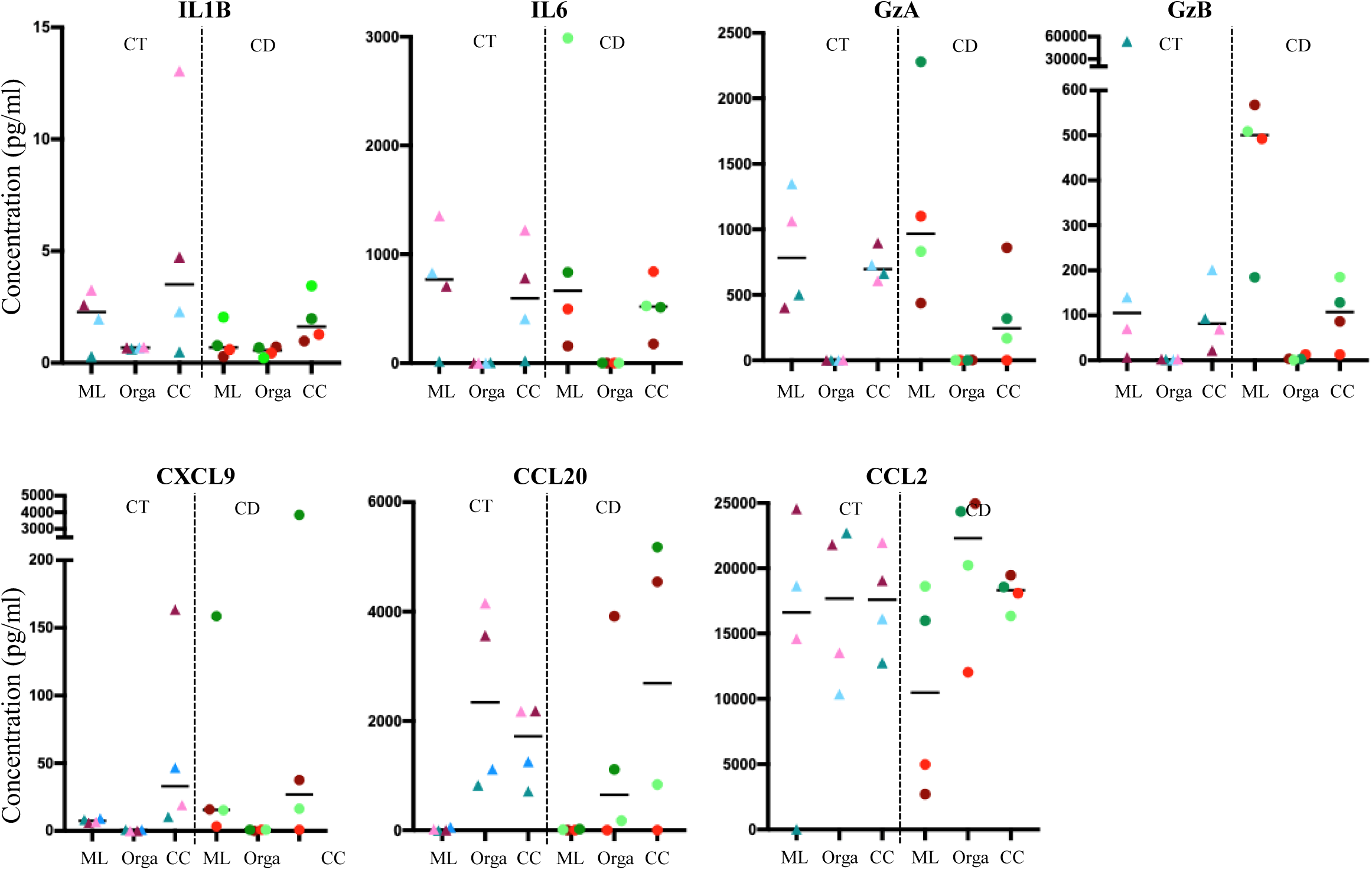
Protein analyze of coculture supernatants. (A) Quantification of 7 cytokines and chemokines (IL1B, IL6, CXCL9, CCL20, CCL2, Granzyme A and B) concentration in the supernatants of 4 control and 4 CD patients cocultures All experiments were made in GM; CT: control patients; CD: CD patients; CC: coculture, Orga: organoids alone, ML: mucosal lymphocytes alone.

## Notes

### Competing Interest Statement

The authors have declared no competing interest.

